# Splice sites obey the power-law during splicing in leukemia cells

**DOI:** 10.1101/2021.05.24.445432

**Authors:** Vasily Grinev, Natalia Siomava, Laurent Vallar, Petr Nazarov

**Affiliations:** Department of Genetics, Faculty of Biology, Belarusian State University, 220030 Minsk, Belarus; College of Health Sciences, Abu Dhabi University, Abu Dhabi, UAE; Quantitative Biology Unit and Department of Oncology, Luxembourg Institute of Health, L-1445 Strassen, Luxembourg

## Abstract

Alternative splicing is an essential characteristic of living cells that usually infers a various exon-exon junction governed by different splice sites. The traditional classification based on the mode of use designates splice site to one of the two groups, constitutive or alternative. Here, we considered another criterion and reorganized splice sites into “unisplice” and “multisplice” groups according to the number of undertaken splicing events. This approach provided us with a new insight in the organization and functionality of leukemia cells. We determined features associated with uni- and multisplice sites and found that combinatorics of these sites follows strict rules of the power-law in the t(8;21)-positive leukemia cells. We also found that system splicing characteristics of the transcriptome of leukemia cells remained persistent after drastic changes in the transcript composition caused by knockdown of the RUNX1-RUNX1T1 oncogene. In this work, we show for the first time that leukemia cells possess a sub-set of unisplice sites with a hidden multisplice potential. These findings reveal a new side in organization and functioning of the leukemic cells and open up new perspectives in the study of the t(8;21)-positive leukemia.

## INTRODUCTION

Alternative splicing is a unique process that allows a cell to archive enormous amounts of information with the restricted capacities of the nucleotide sequence and to unzip it when required. Such approach of information encoding significantly expands the complexity and potential possibilities of a genome with limited starting material packed in a limited space of the nucleus. This phenomenon is widespread in various living organisms, including human. It was estimated that about 95% of human genes are able to produce two or more distinct populations of RNA isoforms (1). Alternative transcript isoforms may be generated by a single cell or by populations of the same cell type (1–2). Additionally, different tissue of the same individual or the same tissue of different individuals (1, 4) may produce different transcripts at different stages of human development (5–6).

Alternative splicing significantly expands the complexity of a transcriptome with limited starting material. Through this process, a single gene can produce a wide variety of RNA molecules. Some of these molecules encode proteins of different structure and function (7), while others are truncated and non-coding. The latter usually play regulatory roles in various biological processes (8). Altogether, functional and regulatory RNA molecules of the same gene form a sub-network, which is tightly integrated into the global regulatory network of a normal or mutated cell, providing flexibility in adaptation and functioning (9–10).

In order to produce different RNA isoforms each multi-exon human gene contains constitutive and alternative splice sites. These sites are specifically distributed along the gene body and flank respective exons assigning them to one of the two groups: constitutive and alternative. In fact, various splice sites of a mature RNA are mount points that are under selection and control of different cis- and trans-factors. A significant progress in the identification of such regulatory elements has been achieved in recent years. In particular, various factors were discovered to selectively control either constitutive or alternative splice sites and exons (11) and complex interaction networks were elaborated in order to explain coordination of these factors in the cell (12–13).

Unfortunately, the traditional classification of splice sites and exons into constitutive and alternative does not fulfil all necessities of the modern research. In this paper, we use an alternative criterion grouping sites and exons by the number of undertaken splicing events. We distinguish “unisplice” sites and exons that undergo one unique splicing event and “multisplice” sites and exons that participate in two or more different splicing events. In this classification, the multisplice sites and exons are specifically of interest because they represent crucial hub points of the splicing process. Importantly, unification of constitutive and alternative splice sites into the multisplice group, does not simply imply unification of their particular properties. In contrast, multisplice sites and exons possess unique properties that often underestimated and remain unclear.

In our recent work, we used the RUNX1-RUNX1T1 fusion gene, present in the t(8;21)-positive acute myeloid leukemia (AML) cells, to show that distribution of exons flanked by constitutive and alternative splice sites follows the power-law (14). In that work, we identified features (potential determinants) associated with the power-law behavior of RUNX1-RUNX1T1 exons during splicing. This finding is of great interest because it shows that alternative splicing of RUNX1–RUNX1T1 follows strict rules of the local combinatorics. The discovered splicing pattern was testes in a complex but single gene. Therefore, we do not know whether that discovery was gene specific or not and whether the proposed model may be generalized and brought to a broader context. We address this question in this paper and present an extended analysis of the whole transcriptome of t(8;21)-positive leukemia cells along with the public data deposited in GenBank. For this, we used nascent and total RNA isolated from the t(8;21)-positive AML model cell line Kasumi-1 with unaltered or siRNA-mediated down-regulated expression of the RUNX1-RUNX1T1 fusion gene.

We show that trends observed in the alternative splicing of RUNX1-RUNX1T1 hold true for the entire cell line, viz., the usage of different splice sites and exons follows the power-law behaviour at the level of nascent and total RNA. Surprisingly, system splicing characteristics of the Kasumi-1 transcriptome remained unaltered despite the dramatic qualitative and quantitative reorganization of the transcript composition and functional changes after knockdown of the RUNX1-RUNX1T1 gene expression. The overall shape of the splice sites and exon distribution did not change and, therefore, can be considered as invariant in this cell type. Using the alternative classification, we analyzed uni- and multisplice sites and described specific features associated with them, the most important of which are the conservatism of adjacent genetic elements, distances to epigenetics marks, and short sequence motifs. Finally, we identified a sub-set of unisplice sites with hidden multisplice potential.

These findings clearly illustrate that combinatorics of splice sites and exons is guided by strict rules that can be in common for single genes and the whole cells. Discovery of the same multi-level behaviour may possibly indicate a uniformity of the power-low model in leukemic or even normal healthy cells. This finding opens new perspectives in the study of the t(8;21)-positive form of AML and provides new insights in the organization and functionality of leukemic cells.

## MATHERIALS AND METHODS

A detailed description of the experimental procedures and all steps of our analytical pipeline are presented in Supplementary Data. A brief description of the most important steps is given below.

### Experimental and public data

This study is based on genome-wide data obtained from mismatch siRNA (siMM) and anti-RUNX1-RUNX1T1 siRNA (siRR) treated Kasumi-1 cells. All microarray, RNA-seq, ChIP-seq, and DNase I hypersensitivity sites data used for this study were previously described (15–18). All raw data are freely available in GEO, SRA or ENA under the studies PRJNA142965, PRJNA236604 and PRJNA674515. In addition to this data, the whole set of GenBank human mRNA and ESTs sequences and genomic coordinates of the human CpGs islands were downloaded via UCSC Genome Browser (19). Finally, annotated transcriptional models of the human genes were retrieved from different sources listed in Supplementary Table S2.

### RNA-seq reads alignment, splicing events identification, and full-length RNA assembling

A global alignment of RNA-seq reads against the GRCh38/hg38 human reference genome and identification of splicing events was carried out according to the protocol by Liao Y. et al. (20). For each RNA sample, a parsimonious set of transcripts was assembled with Cufflinks (21). Individual Cufflinks assembled transcriptomes were merged into one consolidated set of transcripts with Cuffmerge (22). This consolidated set of transcripts was submitted to Cuffdiff (23) for the simultaneous calculation of the transcript abundance and differential expression and finally filtered against i) unstranded transcripts, ii) too short transcripts (<300 nucleotides), iii) transcripts with too short exon(-s) (<25 nucleotides), iv) transcripts with too short intron(-s) (<50 nucleotides), and vi) transcripts with low abundance (fragments per kilobase of transcript per million mapped reads, or FPKM, below 1).

### Reconstruction of splicing graphs and calculation of graph topology

In this paper, we used two different concepts of splicing graphs. First, we reconstructed exon graphs from our Cufflinks assemblies or public sequences. Each exon graph is a set of exons (vertices or nodes of graph) connected to each other via set of splicing events (edges or links of graph) (24). Second, we also reconstructed splice sites graphs from mapped RNA-seq reads or alternatively from our Cufflinks assemblies or public sequences. Each splice sites graph is a set of splice sites as vertices and the connecting edges represent the intermediate exons and introns (25–27). The graphical representation of both concepts of splicing graph is given in the Supplementary Figure S1. For reconstructed graphs, we calculated three variants of the vertices degree as the number of ingoing, outgoing, or all adjacent edges. In the splicing graph theory, the splicing degree of a vertex of a splicing graph is the number of alternative splicing events involving a given splice site or exon. Additionally, for each splicing graph, biologically relevant global and local topological indices were calculated with the respective functions from R/Bioconductor library igraph (28).

### Fitting of statistical models to empirical data

Seven closest statistical models were selected for further fitting to empirical data: power-law, power-law with exponential cut-off, exponential, stretched exponential (or complementary cumulative Weibull), Yule-Simon, log-normal and Poisson. We fitted selected statistical models to empirical data according to the “x_min_” paradigm (29). Finally, goodness-of-fit test, log-likelihood ratio test, Kolmogorov-Smirnov test, and Akaike and Bayesian information criteria were used to assess the plausibility of the statistical hypothesis and for the direct comparison of alternative statistical models (29–31).

### Listing of features associated with splice sites

Every splice site was annotated with sequence, sequence-related, functional, and structural features that were extracted from four types of genomic/RNA elements: 100-bp fragment of the upstream exon (USE), 300-bp fragment from the 5’ end of the intron (USIF), 300-bp fragment from the 3’ end of the intron (DSIF) and 100-bp fragment of the downstream exon (DSE). Additionally, each splice site was described with a set of the nearest epigenetic markers. In total, the list of features included 1680 items.

### Data mining with the random forest meta-classifier

Our primary data matrix included splicing degrees of splice sites as a dependent response variable and all features associated with splice sites were interpreted as independent predictors of the response variable. This matrix was filtered against non-informative features (32) and used for machine learning. Machine learning was carried out with the package randomForest in a classification forests’ mode (33–34). The importance of each feature was determined via calculation of the total decrease in the node impurities from splitting on the feature averaged over all classification trees in random forest. We also applied a recursive algorithm with five-fold cross-validation to select the minimal set of important features required for the accurate classification.

## RESULTS

### Power-law behaviour of different splice sites and exons in alternative splicing

This part of work was based on RNA-seq data obtained from total RNA samples of the t(8;21)-positive AML cell line Kasumi-1 (Supplementary Materials). With these RNA data, we created a transcriptome assembly and identified all splicing events taking place in the Kasumi-1 transcriptome (Supplementary Methods). To formalize the following analysis, we converted a matrix of identified splicing events and assembled transcripts into splice sites and exon graphs, respectively (Supplementary Methods). We applied the idea of the vertex degree from the graph theory (35) to quantify the usage of individual splice sites or exons in various splicing events. In the splicing graph theory, the splicing degree of a vertex is a number of alternative splicing events involving the given splice site or exon (24, 36) (see Supplementary Methods and Supplementary Figure 1 for future explanations).

We found that splicing degrees of splice sites and exons followed a highly asymmetric right-tailed distribution. The vast majority of these elements had low degree, while a small set of splice sites and exons was characterized with high values (Figure 1). Distributions with right tails form a broad statistical family and selection of the right statistical model for this kind of empirical data is not a trivial task (37). Meanwhile, an accurate estimation of the distribution shape allows us to i) describe the distribution properly, ii) formulate a reasonable hypothesis about the mechanism that generates the distribution and iii) predict properties and capacities of the distribution. Therefore, to find an appropriate statistical model, we used a three-step approach based on the mathematical formalism developed by Clauset A. et al. (29, 37–38).

**Figure 1.**
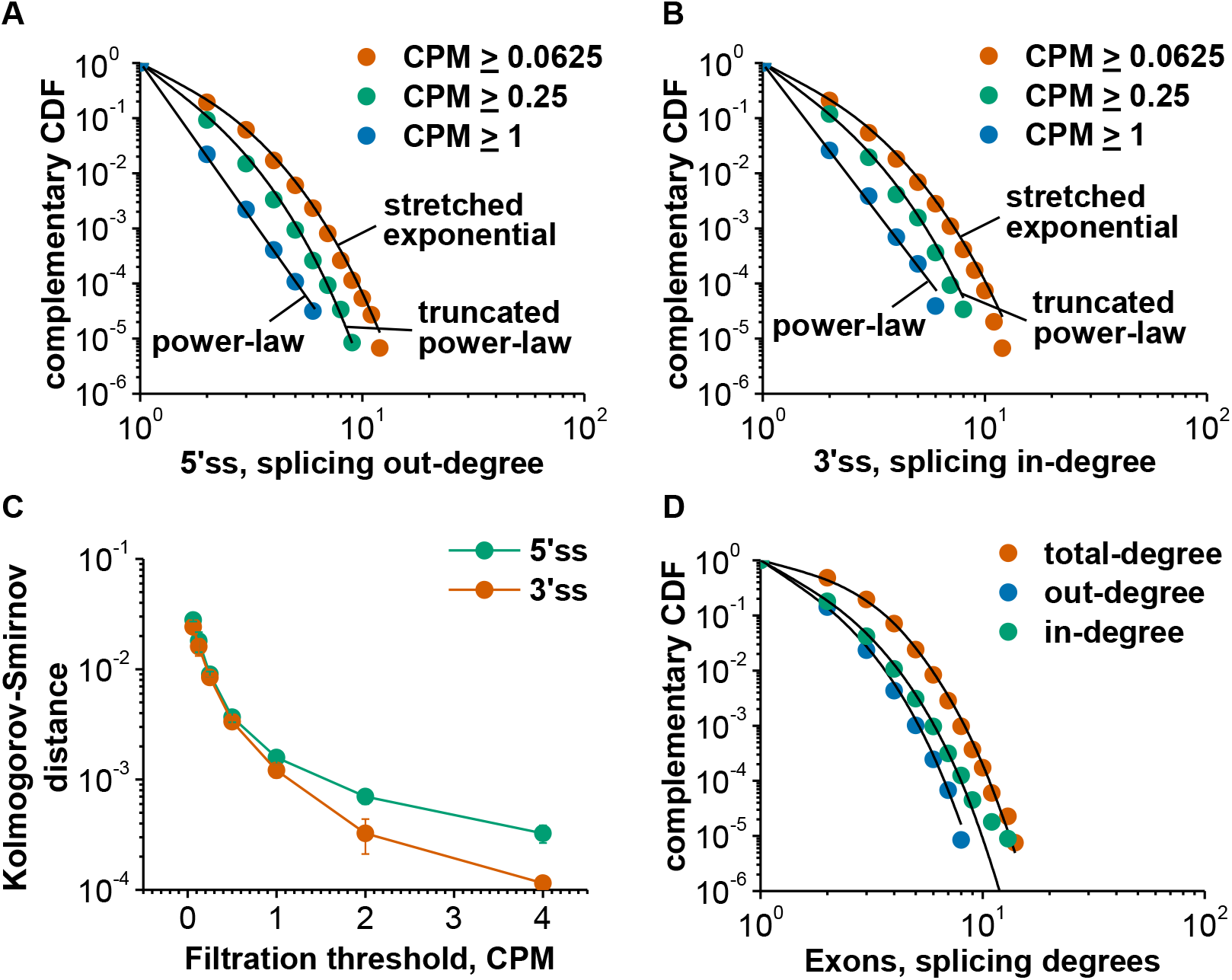
Statistical analysis confirms the power-law component in the transcriptome of Kasumi-1 cells. (A) Log-log plot of the complementary cumulative distribution function (CDF) for the splicing out-degrees of 5’ splice sites at different number of supporting reads. (B) Log-log plot of the complementary CDF for the splicing in-degrees of 3’ splice sites at different number of supporting reads. For each dataset in (A) and (B), the best fits of the most plausible statistical model are shown. (C) The power-law model can be fitted with high accuracy to empirical data after removing poorly supported splicing events. This plot is based on three independent sequencing datasets of Kasumi-1 transcriptome. Each data point is shown as the mean ± SD. (D) Log-log plot of the complementary CDF for the splicing degrees of exons from Cufflinks assembled transcripts. For this dataset, truncated power-law is the most plausible statistical model for each empirical distribution.

First, we rejected those statistical models that clearly did not fit to our empirical data and chose seven best describing models: power-law, truncated power-law (or power-law with exponential cut-off), Yule-Simon, exponential, stretched exponential (or complementary cumulative Weibull distribution), Poisson, and log-normal. We started from the power-law model and fitted it to the tail of the empirical distribution according to the “x_min_” paradigm (29). We accepted the power-law model as the plausible one if it had covered a significant part of the empirical data and passed the goodness-of-fit test. If not, we followed the conventional approach and analyzed the entire range of data. Finally, log-likelihood ratio test, Kolmogorov-Smirnov statistic, and/or information criteria AIC and BIC were used to assess the plausibility of the statistical hypothesis and to directly compare alternative statistical models (see Supplementary Methods for future details) (29–31).

For splice sites, we observed an exponential component in distributions of splicing degrees obtained from the unfiltered splice sites graph. However, we substantially reduced this component by filtering out poorly supported splicing events. We fitted the power-law model to the empirical data with high accuracy for the reliably detected splicing events (normalized number of supporting reads, or CPM, ≥ 1). Eventually, this model was the most plausible among the other six. This observation was valid for both 5′ and 3′ splice sites (Figure 1A-C).

For exons, there was found a clear truncated power-law distribution of splicing degrees (Figure 1D). This model of distribution was the most plausible among the tested statistical models for in- and out-degrees of exons. As for total-degrees, the log-likelihood ratio test gave a preference for two models: the truncated power-law model and the stretched exponential model. The applied test did not favor one model over the other, while Kolmogorov-Smirnov statistic and information criteria highlighted a slight preference of the truncated power-law model. In contrast to the results of splice sites, we observed only a little change in shape of the exon splicing degree distribution after filtering assembled transcripts against low abundance (data not shown). These results indicated that behavior of splice sites and exons during splicing followed a power-law in t(8;21) AML cells and the power-law may be present along with the exponential component.

In our RNA datasets, the maximum splicing degree was 21 (or less) (Figure 2C), while typical data with the power-law distribution cover several orders of magnitude (29, 37–38). It is well known that only a certain part of the overall gene potential to alternative splicing can be realized in one cell type. We assumed that it might account for the low maximum values of degrees in our datasets. To test this hypothesis, we extended our analysis to public data deposited in GenBank and to human gene models generated by different manually curated or fully automatic annotation systems (Supplementary Materials). We found that in the whole human transcriptome values of splicing degrees lied in a range of two orders of magnitude (Supplementary Figure S2 and Supplementary Table S2).

**Figure 2.**
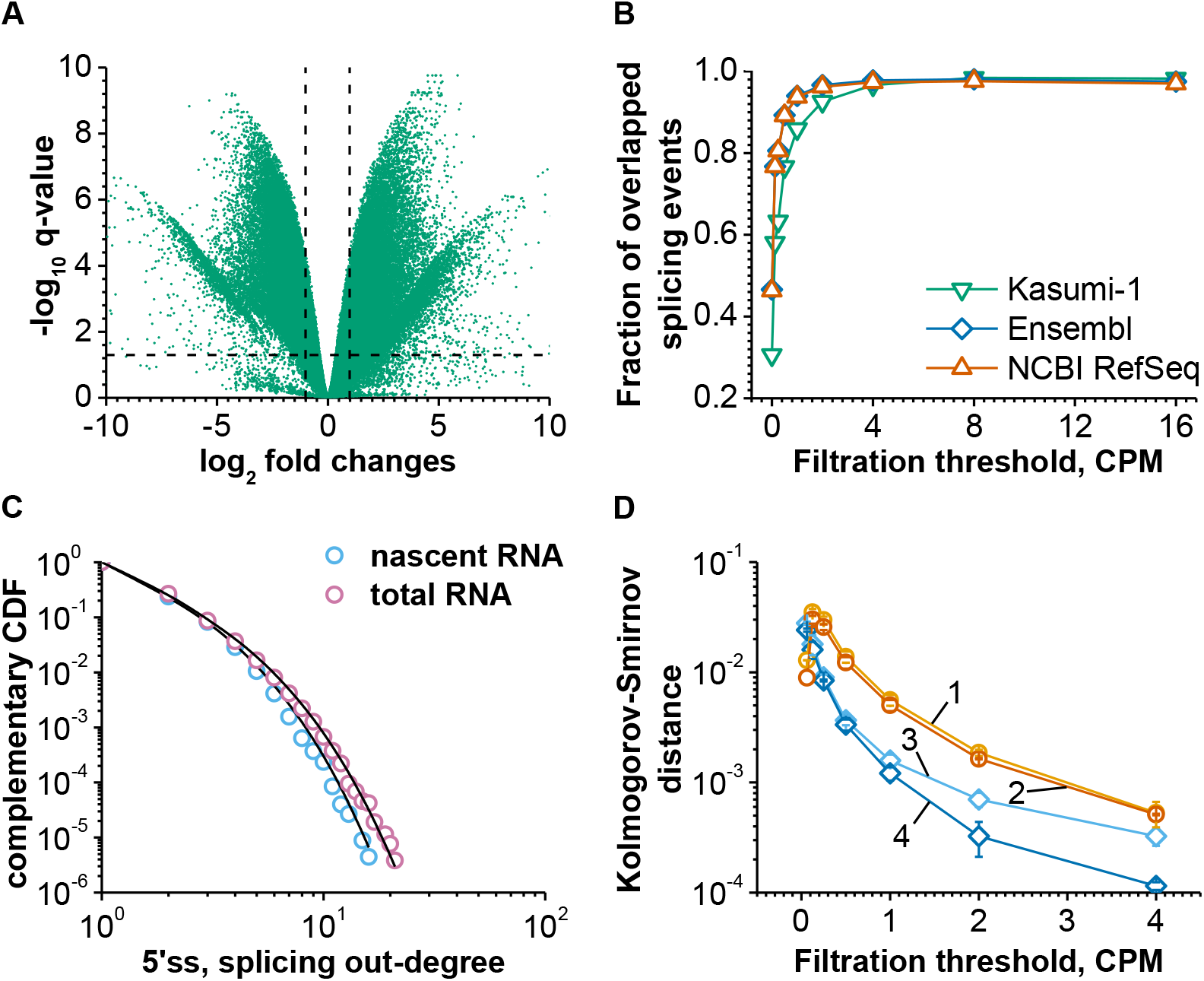
In Kasumi-1 cells, the power-law component can be detected at nascent RNA level of the transcriptome organization. (A) Abundance profiles of splicing events are considerably different between nascent and total RNA fractions. The horizontal dotted line corresponds to the q-value = 0.05. The vertical dotted lines mark the positions of 2-fold differences. (B) A significant overlap between the diversity of splicing events was detected in the nascent RNA fraction and splicing events identified in total RNA of Kasumi-1 cells or total RNA annotated in public databases. Poorly supported splicing events were removed in this plot. (C) Log-log plot of the complementary CDF for the splicing out-degrees of 5’ splice sites from unfiltered nascent and total RNA datasets. For each dataset, the best fit of the stretched exponential model is shown. (D) The power-law model can be fitted to empirical data obtained from nascent RNA with high accuracy after removing poorly supported splicing events. This plot is based on three independent sequencing results of Kasumi-1 transcriptome. Each data point is shown as the mean ± SD. Lines 1 and 2 correspond to splicing out- and in-degrees of 5’ and 3’ splice sites from nascent RNA, respectively. Lines 3 and 4 represent splicing out- and in-degrees of 5’ and 3’ splice sites from total RNA, respectively. Modeling results with total RNA dataset are given for comparison.

### Power-law behavior of splice sites is detected in nascent RNA

In nascent and total RNA fractions of the Kasumi-1 transcriptome, we identified 308329 and 363914 splicing events, respectively. Abundance profiles of these events were considerably different between the two fractions of RNA (Figure 2A). However, there was a significant overlap between the diversity of splicing events detected in nascent and total RNA of Kasumi-1 cells or annotated in public databases, especially after removing the poorly supported splicing events (Figure 2B).

Shape of the distribution of splice site degrees was similar in nascent and total RNA. In particular, we found that in the nascent RNA dataset splice site degrees followed a stretched exponential model without any filtration against poorly supported splicing events was applied. It was identical to the distribution of splice site degrees from the total RNA dataset (Figure 1A-B and Figure 2C).

At the same time, filtration against low values of CPM led to the appearance of the truncated power-law or pure power-law depending on the depth of filtration (Figure 2D). These data indicated that in t(8;21) AML cells the power-law behavior of splice sites during splicing was already present at the nascent RNA level of the transcriptome. However, we needed a stronger depth filtration of the nascent RNA dataset to achieve the same fitting accuracy of the power-law model that we achieved for total RNA (Figure 2D). This was probably due to a larger technical variability of nascent RNA compared to the total RNA dataset (Supplementary Figure S3).

### Power-law behavior of splice sites is invariant in Kasumi-1 transcriptome

It was shown that specific siRNA-mediated knockdown of the RUNX1-RUNX1T1 fusion gene expression leads to large-scale structural and functional changes in the transcriptome of t(8;21) AML cells (16–17, 39–40) and our results supported the finding of that structural changes. In particular, at the gene level, we observed a statistically significant (p < 0.01, q < 0.1) and at least 2-fold difference in expression of 789 genes in the siMM versus siRR treated Kasumi-1 cells (Figure 3A). Similarly, at the transcript level, we identified 1013 transcripts that were differentially expressed in the two states of leukemia cells (Supplementary Figure S4A) and additional 2363 transcripts that were expressed in only one of the two siRNAs treatment conditions. We also found that 95 genes showed significantly different RNA splicing maps at the level of exon-exon junctions in the siMM comparing to siRR treated Kasumi-1 cells (data not shown). Additionally, in the two states of leukemia cells, treated with siMM and siRR, splice site graphs differed by 19417 vertices and 11163 edges and exon graphs differed by 4597 vertices and 5740 edges.

**Figure 3.**
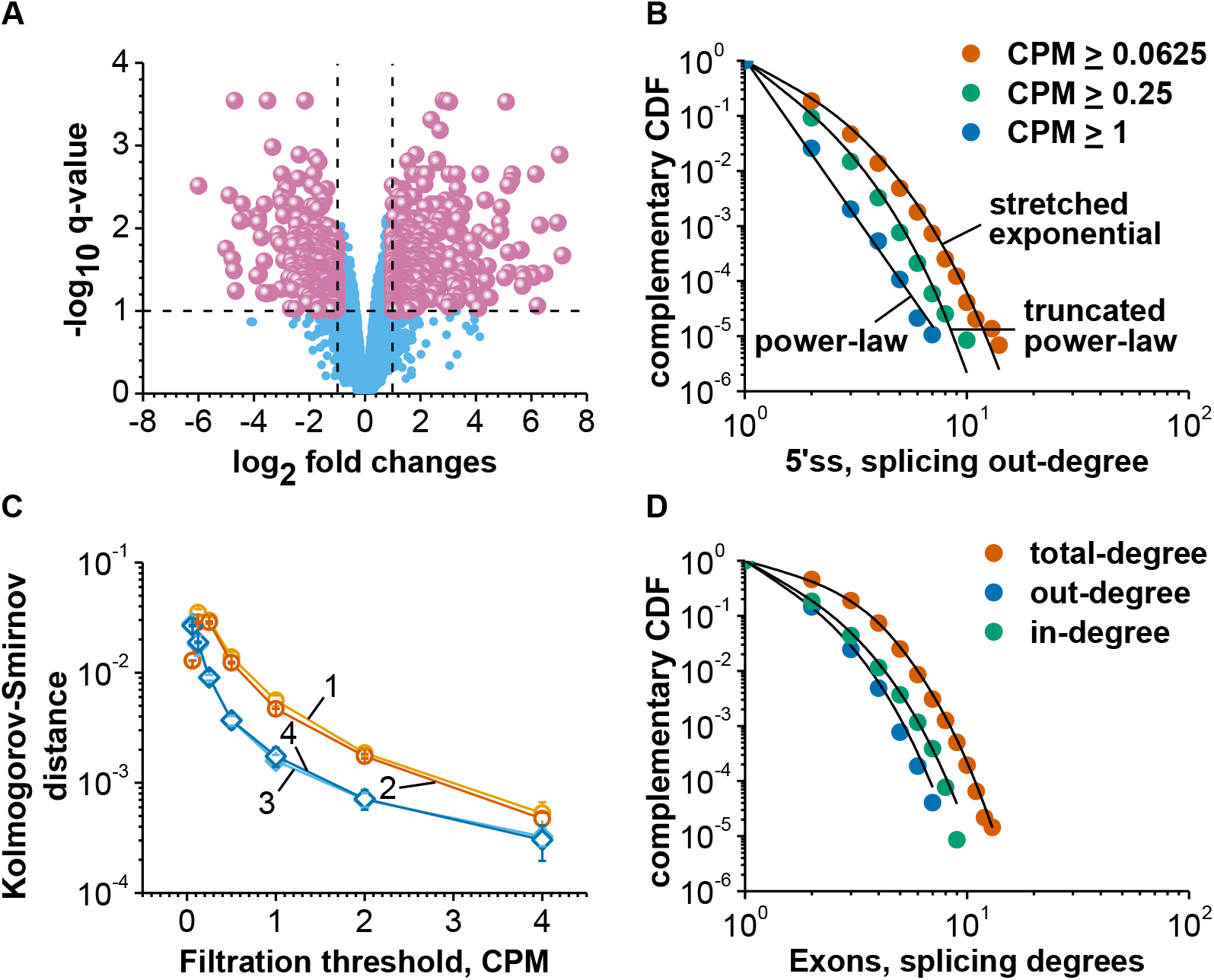
Large-scale changes in the gene expression do not affect shape of distribution of splicing degrees in Kasumi-1 transcriptome. (A) At the level of total RNA, standard limma test supported a significant difference in expression of 794 genes between siMM and siRR treated leukemia cells. The horizontal dotted line corresponds to the q-value = 0.1; vertical dotted lines mark the positions of 2-fold differences. (B) Log-log plot of the complementary CDF for the splicing out-degrees of 5’ splice sites at different numbers of supporting reads. For each dataset, the best fit of the most plausible statistical model is shown. This plot is based on the data obtained from total RNA of siRR treated leukemia cells. (C) The power-law model can be fitted equally well to data obtained from both siMM and siRR treated leukemia cells. In this plot, empirical distributions of the splicing out-degrees of 5’ splice sites were analyzed. Each data point is shown as the mean ± SD from three independent leukemia cells sequencing repeats. Lines 1 and 2 correspond to the nascent RNA dataset; lines 3 and 4 represent the total RNA dataset; lines 1 and 3 show the data from the siMM treated leukemia cells; and lines 2 and 4 correspond to the data from siRR treated leukemia cells. (D) Log-log plot of the complementary CDF for the splicing degrees of exons from Cufflinks assembled transcripts. For this dataset, truncated power-law is the most plausible statistical model for each empirical distribution. The plot is based on the data obtained from total RNA of siRR treated leukemia cells.

Despite drastic changes in the composition of the transcriptome, shape of the distribution of splicing degrees of splice sites was identical in both datasets, obtained from the siMM and siRR treated Kasumi-1 cells (compare Figure 1A and Figure 3B). We also found that for reliably detected splicing events the power-law model was the most plausible one fitting well to the empirical data. These observations were valid for 5′ and 3′ splice sites in both nascent and total RNA datasets (Figure 3B-C, Supplementary Figure S4B). At the exon level, there was a clear truncated power-law distribution of exon splicing degrees in the dataset obtained from anti-RUNX1-RUNX1T1 siRNA treated leukemia cells (Figure 3D). Such kind of distribution was identical to the one of exon splicing degrees in the dataset from the mismatch siRNA treated Kasumi-1 cells (compare Figure 3D with Figure 1D).

Finally, we compared the two analyzed states of the leukemic transcriptome with graph invariants. For this, we selected a set of local and global topological indices that can be interpreted from the biological point of view (Supplementary Section 2.13). We applied local indices to consolidated splicing graphs inferred from total RNA datasets for the two different siRNA treatments. Global indices were used to describe the topology of splicing graphs reconstructed from individual RNA samples. With this approach, we did not find systemic differences between the two states of the Kasumi-1 transcriptome (Table 1 and Supplementary Table S3). Thus, distributions of alpha centrality scores and the number of complete events were identical in the mismatch and anti-RUNX1-RUNX1T1 siRNAs treated Kasumi-1 transcriptomes (Supplementary Figure S5). This shows that systemic characteristics of the leukemic transcriptome remained unchanged despite the dramatic qualitative and quantitative reorganization of the transcript composition and functional changes after knockdown of the RUNX1-RUNX1T1 gene expression. Similarly, shape of the distribution of splicing degrees remained the same for both splice sites and exons. Consequently, the observed distribution can be considered as invariant in this type of leukemia cells.

### Identification of features distinguishing uni- and multisplice sites

In each our dataset, we grouped all splice sites in two classes: i) class #1 of unisplice sites with the splicing degree = 1 and ii) class #2 of multisplice sites with the splicing degree > 1. We next described every splice site with a compendium of 1680 sequence, sequence-related, functional, structural, and epigenetic features (Figure 4A, Supplementary Methods 2.14, Supplementary Table S4). We also applied a random forest meta-classifier to find out features significantly associated with each class of splice sites (Supplementary Methods 2.15).

**Figure 4.**
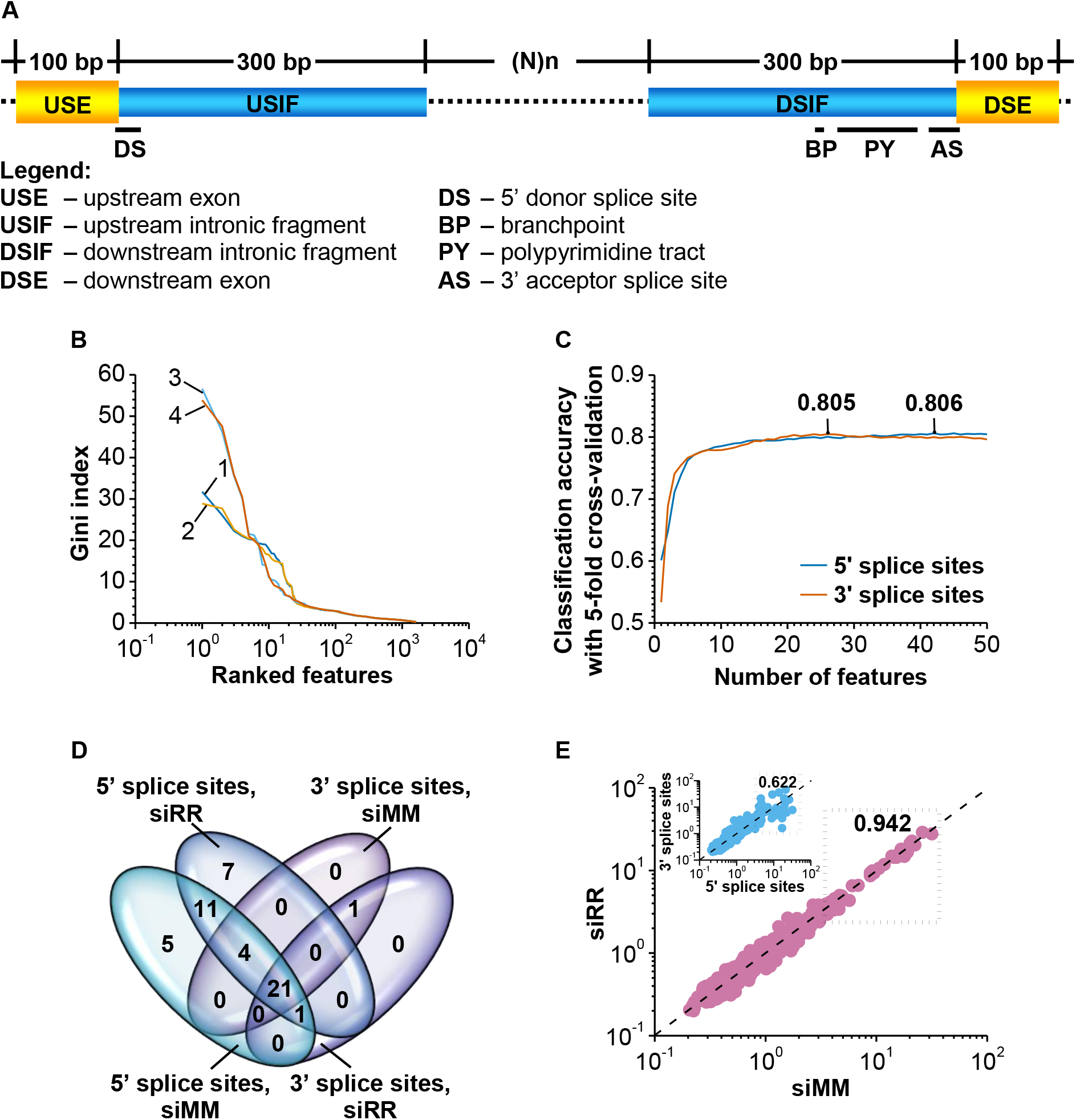
A small set of features distinguishes two classes of splice sites in Kasumi-1 cells. (A) Every splice site was annotated with the following features: sequence, sequence-related, functional, and structural. They all were extracted from four types of genomic/RNA elements: 100-bp fragment of the upstream exon (USE), 300-bp fragment from the 5’ end of the intron (USIF), 300-bp fragment from the 3’ end of the intron (DSIF), and 100-bp fragment of the downstream exon (DSE). Each splice site was additionally described with a set of the nearest epigenetic markers. In total, the complete list of features included 1680 items. (B) Various features are not equivalent in the impact value in determining the class of a splice site. The importance of different features was assessed with Gini index in a random forest-based classification procedure. Lines 1 and 2 show the importance of features in the classification of 5’ splice sites from siMM and siRR treated leukemia cells, respectively. Lines 3 and 4 show the same metrics for 3’ splice sites. (C) Forty-two features of 5’ and 26 of 3’ splice sites features are good enough to properly classify the sites. The numbers on top of lines indicate maximum values of classification accuracy. These results were obtained with a recursive feature selection algorithm and 5-fold cross-validation for data from the siMM treated leukemia cells. (D) Lists of selected features overlap significantly between different types of splice sites and siRNA treatment conditions. (E) The importance of features is irrespective of the state of leukemia cells but it depends on the type of the splice site. Scatterplot represents the importance of various features in determination the 5’ splice site class in siMM and siRR treated leukemia cells. The inner panel shows relationships between the importance of different features in the 5’ and 3’ splice sites determination in the siMM treated leukemia cells. Dotted boxes surround the top 50 features and include Spearman’s rank correlation coefficients for these sub-sets of data.

There was clear evidence that different features were not equivalent in determination to which of the two classes each splice site belongs (Figure 4B). Moreover, results of a recursive feature selection with five-fold cross-validation (Supplementary Methods 2.15) demonstrated that high classification accuracy can be achieved by applying only 50 most important features (Figure 4C). This conclusion was valid for nascent and total RNA from both siMM and siRR treated Kasumi-1 cells. However, 3’ splice sites could be accurately classified with less number of features than 5′ splice sites (Figure 4C and D) and selected features were ranked differently for two types of splice sites (Figure 4B and E).

A short list of selected features included items from all groups of features mentioned above (Supplementary Figure S6). First of all, we found that both the strength of splice sites and the size of clusters of exons to which a splice site and its partner splice site(-s) belonged (see Supplementary Methods 2.14.4 for future details) demonstrated only a moderate importance for the classification accuracy. Therefore, the existence of unisplice and multisplice sites cannot be explained trivially, i.e. by the difference in the strength of splice sites and non-random distribution of constitutive and alternative splice sites along the body of the gene, as could be expected.

Secondly, we identified that the most important group of features was sequence-related. This group included energy of folding of upstream exons and especially conservatism of upstream and downstream exons as well as the downstream fragment from 3’ ends of introns. It is worth mentioning that not only the average score of conservatism was important, but also the range and variability of this parameter among the indicated genetic elements (Supplementary Figure S6).

The second important group included all five epigenetic marks from the primary list of 1680 features (Supplementary Figure S6). In this group, the distance from splice sites to RNA polymerase II peaks was the key factor in classification of both 5’ and 3’ splice sites. The remaining epigenetic features were also highly important, but to a lesser extent than this distance. A special attention should be paid to the distance from splice sites to RUNX1-RUNX1T1 binding peaks. This parameter was significant in assigning 5’ or 3’ splice sites to either uni- or multisplice class in siMM treated Kasumi-1 cells. Surprisingly, this feature was also important for classification of 5’, but not 3’, splice sites in siRR treated Kasumi-1 cells in spite of the significant knockdown in the expression and reduction in the number of binding peaks of the fusion protein.

Finally, we found a clear association between the classification success of 5’ splice sites and AT/CG-rich short motifs (HNRNPF, HNRNPH1, HNRNPK, KHSRP, PCBP2, SFPQ, and TIA1 binding motifs) and Shengdong’s ESEseqs hexamers located in upstream exons. The same association was found for CA/CG-rich short motifs located in intron sequences adjacent to the 5’ splice site and YBX1 binding motifs located downstream the 3’ end of introns. At the same time, all these motifs were not important for the accuracy of 3’ splice site classification with the exception of CG-rich short motifs located in 5’ ends of adjacent introns and TIA1 binding motifs located in upstream exons.

### Splice sites with hidden multisplice potential in t(8;21)-positive AML cells

With the shortlisted features, a random forest meta-classifier achieved a maximum of 80% in the accuracy of splice site classification (Figure 4C). We divided all splice sites in succesfully classified (SC) and misclassified (MC) by iterative run of the meta-classifier. This method converted accuracy in classification of SC splice sites into uni- (SC.1) and multisplice (SC.2) sites to 100% by about 100 iterations (Figure 5A, Supplementary Figure S7 and Supplementary Table S5). Surprisingly, the classification of MC splice sites itself on “unisplice” (MC.1) and “multisplice” (MC.2) sites achieved 90% accuracy after about 500 iterations (Figure 5B, Supplementary Figure S7 and Supplementary Table S5). At the same time, pooling together the opposite classes of splice sites from SC and MC sub-sets led to a significant decrease in the accuracy (Figure 5C). Moreover, the accuracy dropped dramatically when we used SC as a train set and MC as a test set and vice versa (Figure 5D and Supplementary Table S5). These findings held true for 5’ and 3’ splice sites from both siMM and siRR treated Kasumi-1 cells.

**Figure 5.**
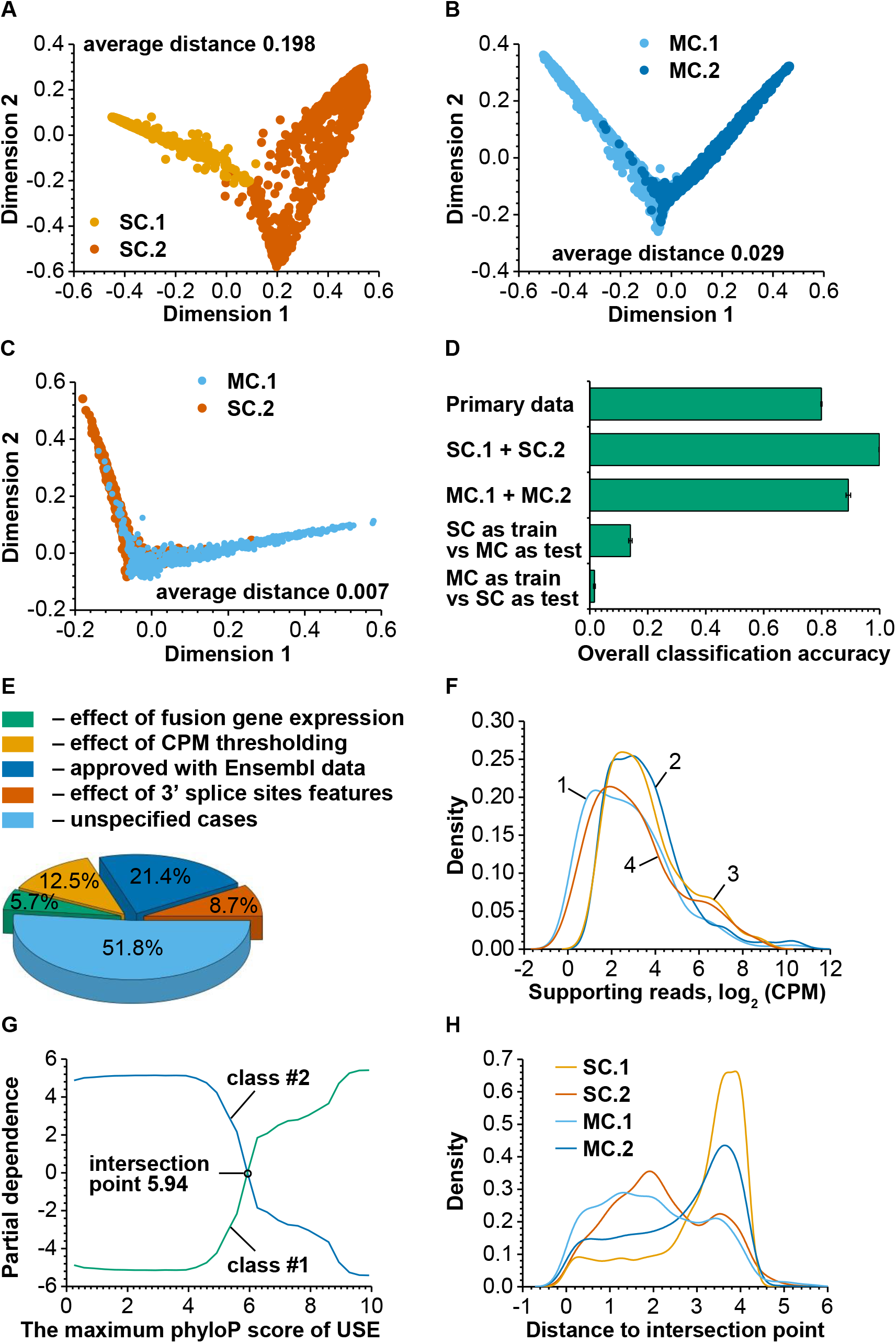
Almost half of misclassified splice sites can be explained. (A-C) Multidimensional scaling of proximity matrices demonstrates a clear difference between two classes of 5’ splice sites in SC (A) and MC (B) sub-sets of splice sites. At the same time, opposite classes of 5’ splice sites from SC and MC sub-sets significantly overlap (C). Plots (A), (B), and (C) are based on the data obtained from the siMM treated Kasumi-1 cells. Average distances in respective proximity matrices are shown. (D) A random forest based accuracy of the 5’ splice site classification. All datasets were obtained from the siMM treated leukemia cells. (E) Distribution of all explained and unexplained cases of misclassified 5’ splice sites from the siMM treated cells. (F) Knockdown of the RUNX1-RUNX1T1 gene in Kasumi-1 cells changes the class of a sub-set of misclassified 5’ splice sites and modulates abundance of these sites in mature RNAs. The x-axis shows the log_2_-transformed normalized number of reads that supports the usage of splice site in the splicing event (-s). Line 1 represents unisplice sites from the siMM treated leukemia cells. Line 2 shows the same splice sites in the siRR treated cells. In last case, the sites changed the class and moved to the multisplice class. Lines 3 and 4 represent multisplice sites in the siMM and RUNX1-RUNX1T1 knockdown cells, respectively. In the siRR treated Kasumi-1 cells, these sites belong to the unisplice class. (G) The effect of different values of the USE maximum phyloP score on the probability of the 5’ splice site class in the siMM treated Kasumi-1 cells dataset. This partial dependence plot gives a graphical depiction of the adjusted effect of a given feature on the class probability in the context of the whole set of other features. The plot is based on results from 100 independent runs of the random forest based meta-classifier and 1000 classification trees per random forest per run. Position of the intersection point and its value are shown. (H) Distances of actual values of the USE maximum phyloP score of 5’ splice sites to the intersection point in the partial dependence plot. This density plot is based on the data from the siMM treated leukemia cells.

The observed classification outcome can only be explained by a complete inversion of splice site classes in the MC sub-set. Namely, features associated with unisplice (or multisplice) sites in the SC sub-set fully corresponded to features of multisplice (or unisplice) sites in the MC sub-set. This hypothesis was further supported by the inversion of partial dependence profiles of features in SC and MC sub-sets (Supplementary Figure S8).

To explain the inversion, we first examined an effect of the RUNX1-RUNX1T1 expression on misclassification of splice sites. We found that knockdown of the RUNX1-RUNX1T1 gene in Kasumi-1 cells changed the class membership of misclassified 5’ splice sites in 5.7-6.1% cases (depending on what dataset was used). This also significantly modulated the abundance of these sites in mature RNAs (Figure 5E and Figure 5F).

Another source of misclassification was linked to incorrect CPM thresholding. In fact, we used a fixed threshold CPM ≥ 1 while preparing the splicing event matrix. A new flexible CPM threshold in the range 0.25 ≥ CPM ≥ 2 could explain additional 12.1-12.7% of misclassified splice sites.

A part of misclassified cases can be also explained by Ensembl data for splicing degrees of splice sites in the whole human transcriptome. These data additionally affected the class membership of 21.4-24% misclassified splice sites. Unfortunately, these data did not provide any mechanistic insight into the observed phenomena in Kasumi-1 cells.

Finally, a part of misclassified cases could be explained by the effect of features of partner splice site (-s). In fact, in our datasets, we described splice sites as pairs of interconnected sites from splicing events. This approach takes into account the influence of features of partner splice site (-s) on the classification outcome for a given splice site. However, in some rare and very specific cases, it may lead to misclassifications because of the strong effect of features of partner splice site (-s). With these considerations in mind, we found that the class membership of additional 5.0-8.9% of misclassified splice sites can be explained by the class membership of their partner splice site (-s).

In total, with the observations mentioned above, we were able to explain about half (43.7-48.2% depending on the dataset) of misclassified cases, while the second half remained unexplained. About 80% of unexplained cases were unisplice sites with features of multisplice sites. The actual values of features of these sites aspired to the intersection point of the partial dependency curves (see Figure 5G and H with the maximum phyloP score of upstream exons as an example). The intersection point was a kind of an equilibrium value of a feature at which no preference can be given to any of the splice site class. It is interesting to note that successfully classified multisplice sites aspired to the intersection point as well, while it remained unclear whether this tendency was present for misclassified unisplice sites (Figure 5H). Thus, the major part of unexplained misclassified unisplice sites had a hidden multisplice potential. We strongly believe that these sites may be reserved for alternative splicing in Kasumi-1 cells.

## DISCUSSION

Traditionally, splice sites (and exons) are subdivided into two major groups: constitutive and alternative, depending on their abundance in mature transcripts of a gene (41–42). This is, however, a very basic classification that sometimes does not cover all needs. In this work, we applied another classification criterion such as the number of splicing events the splice site or exon participates in. Accordingly, we subdivided all splice sites and exons into uni- (one splicing event) and multisplice (more than one splicing event) groups irrespective of their constitutive or alternative status and the alternative splicing mode they are engaged in. This classification allowed us to take a look at the problem of splicing in leukemia cells from a new perspective and to ask new questions.

We found that uni- and multisplice sites involved in splicing process in Kasumi-1 cells followed the pure or truncated power-law. This finding is consistent with our previous observation on behaviour of RUNX1-RUNX1T1 exons flanked by constitutive and alternative splice sites (14). These results are of great importance since a precise definition of the distribution makes it possible to correctly describe data, formulate reasonable hypotheses about mechanism(-s) that drives these data, predict properties and capacities of the object of interest, and improve the quality of future investigations.

At the same time, we have to admit that the reliable approximation of transcriptomic data is an intricate task. The complexity of this task is, in particular, increased by incompleteness of available data, high variability and heteroscedasticity of the data, methodological and/or technical artefacts introduced during the generation, pre-processing, and/or annotation of the data. For instance, working with our Kasumi-1 experimental data, we observed a significant alteration of the distribution shape. It changed from stretched exponential to truncated power-law and to power-law models along with filtration of primary data against the number of reads supporting various splicing events. Using public data, we found an inconsistence between some databases or even between different releases of the same database (data not shown). The approximation was further complicated by the fact that biological data are usually generated under the influence of multiple generative mechanisms, e.g. additive and multiplicative processes. Additionally, asymmetric distributions are known to form a large family with different distributions of similar shape (29, 37). Thus, it is a challenge for many studies to fit experimental results to molecular data (43).

In fact, power-law distributions are mentioned in an enormous variety of different complex systems, from engineering to biological and social sciences (37, 44). Biological systems, as the most complex systems, are particularly rich in this phenomenon, which is manifested at all levels of the life organization, from molecules to ecosystems (44–46). Therefore, it was not a surprise to find splice sites exhibiting the same power-law behaviour. More exciting here is another question: Why is this phenomenon so common in diverse biological systems? We assume that this happens due to the unique properties that the system acquires when it obeys the power-law behavior in its one or several parameters.

The first property of a system with power-law is scale-free (37). The second one is an adaptive flexibility of a system with the power-law component (37, 47–48). The third property of the system with the power-law component is a high sensitivity to targeted attacks and its relative robustness to accidental damages (49–50). We assume that these three properties together permit the splicing process to be flexible and highly adaptive in response to environmental conditions.

In the current study, we created a comprehensive list of features associated with splice sites. We used a recursive selection algorithm with five-fold cross-validation in a random forest mode to find features that are most important for the classification of splice sites. Surprisingly, we found that conservatism of upstream and downstream exons as well as downstream fragments of introns was the most powerful predictor of the splice site class. To obtain an unbiased and reliable outcome at this step, we used two independent conservatism scoring models phyloP and phastCons (51–52). Our finding is in agreement with a very recent observation by Wainberg M. et al. according to which conservation is an unexpectedly powerful indicator of alternative splicing patterns (53).

At the same time and contrary to our expectations, there was found only a limited sub-set of important sequence motifs. This sub-set included manly CG-rich short motifs and binding sites of HNRNPF, HNRNPH1, HNRNPK, and PCBP2 proteins, which are members of ubiquitously expressed heterogeneous nuclear ribonucleoprotein subfamily. The first two ribonucleoproteins were shown to specifically bind G-tracts in pre-mRNAs and regulate alternative splicing (54–55). Interestingly, G-tracts buffer genetic variations of 5’ splice sites acting as evolutionary capacitors of splicing changes (56). This is in agreement with our results highlighting importance of the genetic element conservatism in our classification model.

The other two proteins, HNRNPK and PCBP2, possess a very specific binding preference that distinguishes them from other heterogeneous nuclear ribonucleoproteins. They tenaciously bind to poly(C) repeats, especially to those with the polypyrimidine tract (57–58). A similar polypyrimidine tract binding activity characterizes the SFPQ protein (59), which was also present among other important proteins selected in our pipeline. We also found a strong association of YBX1 motif with the class membership of splice sites. This motif is known to activate splicing by facilitating the recruitment of U2AF65 to weak polypyrimidine tracts through direct protein-protein interactions (60). It was suggested that poly(C) binding proteins play a sufficient role in development of some type of cancer, including leukemia, but the exact mechanisms are not completely clear yet (61–64). Our results contribute to this suggestion and confirm an important role of these proteins as splicing regulators in leukemia cells.

Another association we found was the one between AT-rich short motifs, AT-rich binding proteins (e.g. KHSRP and TIA1), and the classification accuracy. There are limited data available on the splicing regulatory activity of these proteins (65–66). So far, there also has not been found a strong support to their contribution to the leukemia development (67). Indeed, our results provide first evidences that KHSRP and TIA1 proteins may significantly regulate 5’ splice site behaviour in leukemia cells.

Our analysis revealed a significant role of epigenetic marks in assigning a splice site to the one of two classes. Among these predictors, the most interesting feature was a distance from splice sites to RUNX1-RUNX1T1 protein binding peaks. We found that multisplice sites tend to be closer to these peaks. There was a clear interrelation between the sub-set of misclassified splicing states of splice sites and the expression status of the fusion oncogene. This is the first ever evidence that the RUNX1-RUNX1T1 oncogene may affect alternative splicing in t(8;21)-positive AML cells. This influence may happen due to the chromatin remodeling and changes in the transcription control caused by the RUNX1-RUNX1T1 protein (16–17). This is, however, only a preliminary discovery that has to be further tested and evaluated in additional experiments.

In conclusion, we would like to mention here for the first time that the combinatorics of splice sites is follow a strict rule during splicing process in t(8;21)-positive AML cells. This strict behaviour of splice sites is robust to transcriptome perturbations and associated with specific sequence, sequence-related, and epigenetics features. These findings provides a new insight in the organization and functionality of leukemic cells and opens new perspectives in the study of the t(8;21)-positive form of AML. In particular, it introduces new attractive objectives for further investigations, such as the functional role of splice sites with hidden multisplice potential, the exact mechanism(-s) of the influence of conservative genetic elements on behaviour of splice sites, and the splicing regulatory activity of the RUNX1-RUNX1T1 fusion protein.

## SUPPLEMENTARY DATA

Supplementary Data are available online.

## ACKNOWLEDGEMENTS

The authors appreciate a critical review of the manuscript by Dr. Ilia Ilyushonak (Belarusian State University).

## FIGURE LEGENDS

**Table.**
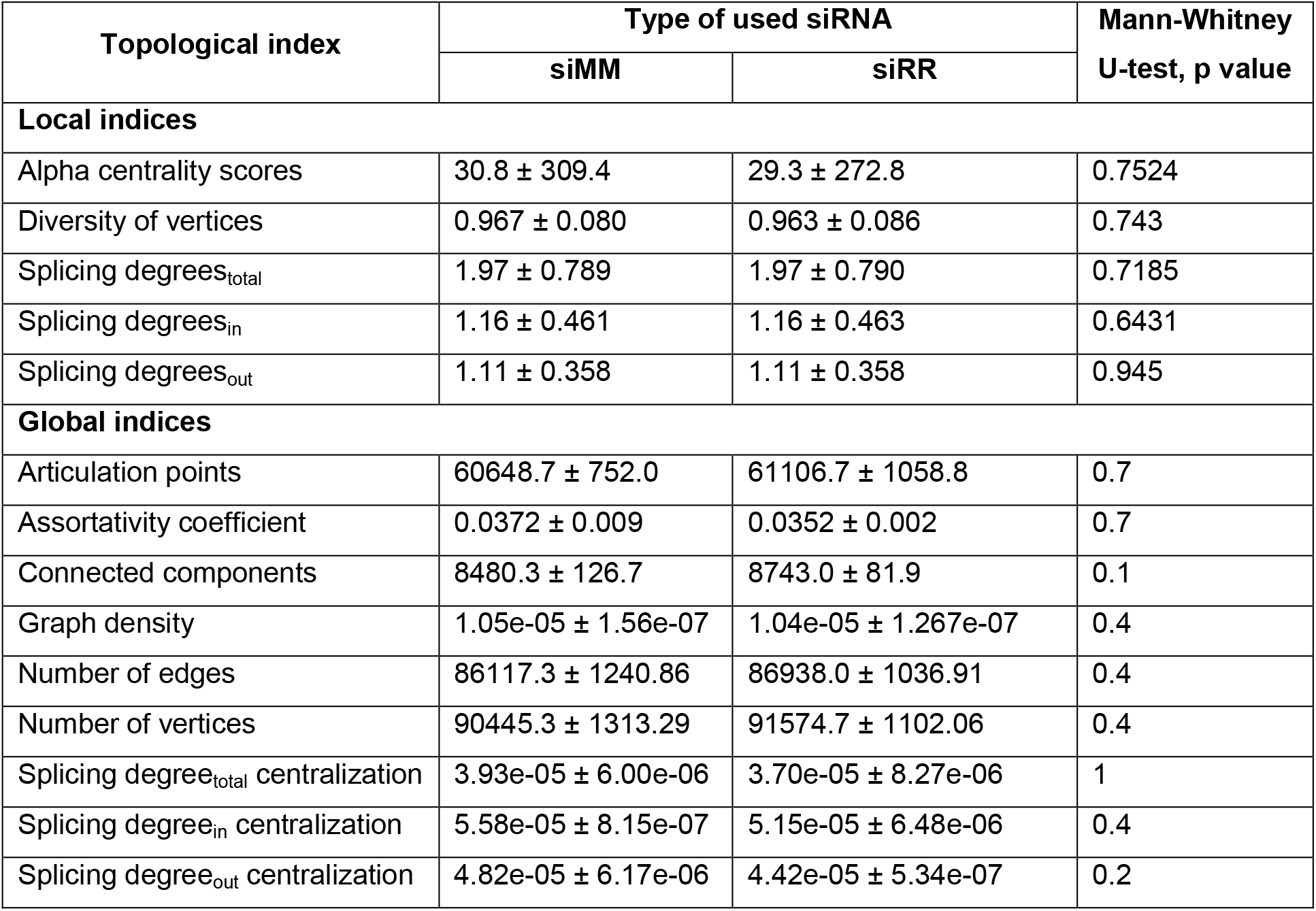
Knockdown of the RUNX1-RUNX1T1 fusion gene expression in Kasumi-1 cells did not lead to any change of the whole transcriptome based exon graphs topology.

| Topological index | Type of used siRNA |  | Mann-Whitney<br>U-test, p value |
| --- | --- | --- | --- |
|  | siMM | siRR |  |
| Local indices |  |  |  |
| Alpha centrality scores | 30.8 ± 309.4 | 29.3 ± 272.8 | 0.7524 |
| Diversity of vertices | 0.967 ± 0.080 | 0.963 ± 0.086 | 0.743 |
| Splicing degrees<sub>total</sub> | 1.97 ± 0.789 | 1.97 ± 0.790 | 0.7185 |
| Splicing degrees<sub>in</sub> | 1.16 ± 0.461 | 1.16 ± 0.463 | 0.6431 |
| Splicing degrees<sub>out</sub> | 1.11 ± 0.358 | 1.11 ± 0.358 | 0.945 |
| Global indices |  |  |  |
| Articulation points | 60648.7 ± 752.0 | 61106.7 ± 1058.8 | 0.7 |
| Assortativity coefficient | 0.0372 ± 0.009 | 0.0352 ± 0.002 | 0.7 |
| Connected components | 8480.3 ± 126.7 | 8743.0 ± 81.9 | 0.1 |
| Graph density | 1.05e-05 ± 1.56e-07 | 1.04e-05 ± 1.267e-07 | 0.4 |
| Number of edges | 86117.3 ± 1240.86 | 86938.0 ± 1036.91 | 0.4 |
| Number of vertices | 90445.3 ± 1313.29 | 91574.7 ± 1102.06 | 0.4 |
| Splicing degree<sub>total</sub> centralization | 3.93e-05 ± 6.00e-06 | 3.70e-05 ± 8.27e-06 | 1 |
| Splicing degree<sub>in</sub> centralization | 5.58e-05 ± 8.15e-07 | 5.15e-05 ± 6.48e-06 | 0.4 |
| Splicing degree<sub>out</sub> centralization | 4.82e-05 ± 6.17e-06 | 4.42e-05 ± 5.34e-07 | 0.2 |

