## Supplementary Data for "Splice sites obey the power-law during splicing in leukemia cells"

### 1. SUPPLEMENTARY MATERIALS

#### 1.1. NGS data

##### *1.1.1. RNA-seq data*

In this study, samples of total RNA from the mismatch or anti-RUNX1-RUNX1T1 siRNA treated Kasumi-1 cells were analysed. Procedures for total and nascent RNA isolation, preparation of libraries, and massively parallel sequencing were previously described (1-3). FASTQ files with raw data are freely available in Sequence Read Archive or European Nucleotide Archive under the studies PRJNA236604 and PRJNA674515.

##### *1.1.2. DNase I hypersensitivity sites data*

Distribution of the DNase I hypersensitivity sites in the genome of Kasumi-1 cells was previously established (4). Raw data are freely available in Sequence Read Archive or European Nucleotide Archive under the study PRJNA142965. Processed data can be downloaded from GEO repository under the sample accession number GSM722713.

##### *1.1.3. ChIP-seq data*

Positions of the H3K9Ac, RNA polymerase II, and RUNX1-RUNX1T1 peaks in the genome of the mismatch and anti-RUNX1-RUNX1T1 siRNA treated Kasumi-1 cells were previously mapped (4). Raw data are freely available in Sequence Read Archive or European Nucleotide Archive under the study PRJNA142965. Processed data can be downloaded from GEO repository under the series accession number GSE29222.

#### 1.2. Public data

##### *1.2.1. Human mRNA and ESTs sequences from GenBank*

The whole set of GenBank human mRNA and ESTs sequences was downloaded via UCSC Genome Browser (5) in GTF format. We used sequences that were pre-aligned against the GRCh38/hg38 reference assembly of the human genome with BLAT. Any sequences with mismatches and less than four aligned blocks were removed. Additionally, sequences with the exon length <25 bp and intron length <50 bp were filtered out. For each sequence, we trimmed terminal exons to remove the uncertainty with genomic coordinates of these exons.

##### *1.2.2. Transcriptional models of human genes*

Annotated transcriptional models of human genes were retrieved from AceView (6), ECgene (7), Ensembl (8), GENCODE (9), NCBI RefSeq (10), and VEGA (11) in GTF format. Download releases of these databases are listed in Supplementary Table S2.

##### *1.2.3. CpGs islands*

Genomic coordinates of the CpGs islands in the human genome were downloaded via FTP from UCSC Genome Browser (4).

#### 2. SUPPLEMENTARY METHODS

##### 2.1. Alignment of RNA-seq reads against the reference genome

GRCh38/hg38 reference assembly of the human genome was downloaded as twoBit file from FTP server of UCSC Genome Browser. It was converted to a standard FASTA format with twoBitToFa utility (4). A hash table for the reference genome was built with the function *buildindex* from R/Bioconductor library Rsubread (12). At this step, 16-mers subreads were extracted in every three bases from the reference genome. The threshold value 24 was used to exclude highly repetitive subreads from the created hash table.

A global alignment of RNA-seq reads against the reference genome was carried out with the function *subjunc* from R/Bioconductor library Rsubread (12) and the created hash table. This function implements a seed-and-vote mapping paradigm for a fast and accurate alignment (12). At this step, we used default settings of the function *subjunc* and collected only uniquely mapped reads with the maximum of 3 mismatched bases in the alignment. The resulting BAM files were sorted with the function *sortBam* and indexed with the function *indexBam*, from R/Bioconductor library Rsamtools (13). Alignment statistics for all our RNA-seq libraries was calculated and presented in Supplementary Table S1.

##### 2.2. Identification and pre-processing of exon-exon junctions

All possible variants of exon-exon junctions were identified according to Liao Y. et al. (12). The resulting BED files were parsed and converted into the primary count matrix of splicing events with an in-laboratory developed R code. This matrix included a full list of identified splicing events with the respective genomic coordinates and number of reads supported each event in every RNA sample.

The primary count matrix was wrapped in a DGEList object (14) and subjected to a multi-step normalization procedure. First, RNA samples were divided into four groups according to the type of RNA and expression status of RUNX1-RUNX1T1. From this, a contrasts parametrization design matrix was developed (14). Second, to calculate an effective size of each RNA-seq library, the scaling factors were estimated using the “trimmed mean of M-values” normalization method (15). Third, the voom normalisation was applied to count data with some degree of heteroscedasticity (16). Fourth, by applying the calculated scaling factors, the voom normalised counts were converted into counts per million (CPM) without the $\log_2$ transformation. Finally, for the fast retrieving of the data and easy subsequent manipulations, the normalized matrix was wrapped in a GRanges object (17) and used in the downstream analytical pipelines.

##### 2.3. Cufflinks-based assembly of transcriptomes

For each sample of total RNA, we used Cufflinks (18) and the respective *subjunc*-generated BAM file to assemble the alignments into a parsimonious set of transcripts and to estimate the relative abundances of these transcripts. Herewith, Cufflinks was not supplied with the reference annotations and a minimal isoform fraction threshold was assigned to 0.05.

Individual Cufflinks assemblies were filtered against i) unstranded transcripts, ii) too short transcripts (<300 nucleotides), iii) transcripts with too short exon(-s) (<25 nucleotides), iv) transcripts with too short intron(-s) (<50 nucleotides), and vi) transcripts with low abundance (fragments per kilobase of transcript per million mapped reads, or FPKM, below 1). Finally, for the fast retrieving of the data and easy subsequent manipulations, the filtered assemblies were converted into objects of the class TranscriptDb (17) and saved as local SQLite databases.

Alternatively, individual Cufflinks assembled transcriptomes were merged into one consolidated set of transcripts with Cuffmerge (19). This set of transcripts was filtered in the way described above but without the last filtration step. The consolidated set of transcripts was initially submitted to Cuffdiff for the simultaneous calculation of the transcript abundance and differential expression (20). Cuffdiff was run in a default mode with the exception of the minimal isoform fraction threshold that was assigned to 0.05. Next, the Cufflinks/Cuffmerge consolidated set was additionally annotated with the Cuffdiff FPKM tracking output data and filtered against transcripts with low abundance. We used FPKM ≥ 1 for at least one siRNA treatment condition as a threshold during this filtration step. Finally, we divided the consolidated filtered annotated set of transcripts into two subsets according to the expression status of transcripts in the siMM and siRR treated Kasumi-1 cells. These subsets were converted into objects of the class TranscriptDb and saved as local SQLite databases.

##### 2.4. Reconstruction of splicing graphs

The concept of the splicing graph was proposed and formalized by Heber S. et al. in 2002 (21). It was a model alternative to linear transcriptional models of structural and functional organization of eukaryotic genes. Later, it was successfully used to solve a number of issues in RNA biology (22-28).

According to the basic concept, splicing graph $G = (V, E)$ can be defined as follows. Let $\{t_1, t_2, t_3, \dots t_n\}$ be a set of RNA transcripts of the gene of interest. Herewith, let each $i^{th}$ transcript be a linearly ordered set of transcribed and spliced subset $V_k$ of genomic positions (in this subset, each $V_l \neq V_m$ for $l \neq m$). Define $V := U_i V_i$ as a set of all transcribed and spliced genomic positions, or exons. Additionally, define $E := U_i E_i$ as a set of all elementary splicing events. $E_i$ led to formation of the complete set of RNA transcripts of the gene of interest. In this context, splicing graph is a set $V$ of exons (vertices or nodes of graph) connected to each other via the set $E$ of splicing events (edges or links of graph). Such a graph is a directed acyclic graph in the sense that exons present in any mature transcript are retained in the correct 5’ to 3’ linear order and forward back edges are prohibited. This classical type of a splicing graph, we called the exon graph.

To reconstruct the exon graph from Cufflinks assembled transcripts or publicly available datasets, each exon was given a unique name corresponding to its genomic coordinates and each assembled transcript was represented as a unique vector of such exons. Every unique vector was transformed into a list of pairwise spliced exons (list of splicing events) according to their linear order in the transcript, beginning with the 5’-terminal exon and ending with the 3’-terminal exon. All lists of splicing events were merged, any duplicates were removed and the consolidated list of splicing events was transformed into a graph object of R/Bioconductor library igraph (29) for subsequent manipulations.

Furthermore, we used a slightly modified concept of splicing graph that was independently proposed by Sperisen's and Sugnet's teams in 2004 (30-31) and formalized by Sammeth M. et al. (32-33). Sammeth's splicing graph is a directed acyclic graph similar to the exon graph. However, in contrast to the exon graph, vertices of Sammeth's splicing graph are splice sites and the connecting edges represent the intermediate exons and introns. This type of splicing graph, we called the splice site graph. The graphical representation of both concepts of splicing graph is given in Supplementary Figure S1.

To reconstruct the splice site graph from Cufflinks assembled transcripts or publicly available datasets, we used R/Bioconductor library SplicingGraphs (34). Alternatively, we developed an original R code to reconstruct the splice site graph directly from the normalized matrix of experimentally identified splicing events. In this case, the splice site graph was reconstructed as a set of sub-graphs of interconnected 5' and 3' splice sites without edges between 3' and 5' splice sites. Our approach prevents a significant loss of information during the transcript assembling. At the same time, this approach permits the analysis of the splice site behaviour, but not behavior of entire exons.

**2.5. Calculation of splicing degrees of splice sites and exons**

In graph theory, the degree of a vertex of a graph is the number of edges adjacent to the vertex (35). Herewith, in-, out-, and total-degree of a vertex is the number of ingoing, outgoing or all adjacent edges, respectively. Similarly, in splicing graph theory, the splicing degree of a vertex of a splicing graph is the number of alternative splicing events involving a given splice site or exon. This basic index was calculated with the standard function *degree* from R/Bioconductor library igraph (29). The graphical representation of the splicing degree concept is given in Supplementary Figure S1.

**2.6. Fitting of the power-law model to empirical data**

***2.6.1. Basic functions***

The probability mass function for the discrete power-law is

$$p(x) = Cx^{-\alpha} \qquad \text{(S1)}$$

where *C* is the normalization constant and $\alpha$ is the exponent or scaling parameter (36-37).

The normalization constant *C* can be calculated as follows:

$$C = \frac{1}{\zeta(\alpha, x_{min})}, \qquad \text{(S2)}$$

where $\zeta$ is the generalized or Hurwitz zeta function:

$$\zeta(\alpha, x_{min}) = \sum_{n=0}^{\infty} (n + x_{min})^{-\alpha}. \qquad \text{(S3)}$$

The normalization constant *C* allows us to reduce the function *p(x)* to 1:

$$\sum_{x=x_{min}}^{\infty} Cp(x) = 1. \qquad \text{(S4)}$$

The cumulative distribution function for the discrete power-law is

$$P(x) = 1 - \frac{\zeta(\alpha, x)}{\zeta(\alpha, x_{min})}. \qquad (S5)$$

The complementary cumulative distribution function for the discrete power-law can be expressed as

$$\overline{P}(x) = \frac{\zeta(\alpha, x)}{\zeta(\alpha, x_{min})}. \qquad (S6)$$

***2.6.2. Estimation of the lower bound $x_{min}$ by Kolmogorov-Smirnov statistic***

In fact, more often the power-law applies only for values greater than some minimum $x_{min}$ (lower bound) but not for the full data set. The correct value of this parameter can be calculated by a simple minimization of Kolmogorov-Smirnov distance *D*. The distance *D* (or Kolmogorov-Smirnov statistic) is simply the maximum distance between the cumulative distribution function of the data and the fitted model:

$$D = \sup_x \left| P(x) - P_{emp}(x) \right|, \qquad (S7)$$

where $sup_x$ is the supremum of the set of distances, $P(x)$ is the hypothesized cumulative distribution function, and $P_{emp}(x)$ is the empirical cumulative distribution function (37-38).

In this work, lower bounds in empirical data were determined with the respective code module from R/Bioconductor library poweRlaw (39).

***2.6.3. Determination of the scaling parameter $\alpha$ value by maximum likelihood estimation***

The method of maximum likelihood gives accurate parameter estimation while fitting the model distribution to the empirical data (40-41). If it is assumed that the empirical data follow a discrete power-law distribution, then the scaling parameter $\alpha$ can be determined by the maximum likelihood estimator (36-37):

$$\alpha \cong 1 + n \left[ \sum_{i=1}^{n} \ln \frac{x_i}{x_{min} - \frac{1}{2}} \right]^{-1}, \qquad (S8)$$

where $x_i$ (at $i$ = 1, 2, 3, …, $n$) are the observed values of $x$ such that $x_i \geq x_{min}$.

This estimator gives results accurate to about 1% or better at $x_{min} \geq 6$. Wherein the statistical error of $\alpha$ can be calculated by the equation (36):

$$\sigma = \frac{\alpha - 1}{\sqrt{n}} + O\left(\frac{1}{n}\right). \qquad (S9)$$

However, if there needs more precision (or $x_{min} < 6$), then the scaling parameter $\alpha$ can be estimated by the direct numerical maximization of the likelihood function itself or its logarithm (36):

$$\mathcal{L}(\alpha) = -n\ln\zeta(\alpha, x_{min}) - \alpha \sum_{i=1}^{n} \ln x_i. \qquad (S10)$$

In this case the statistical error of $\alpha$ can be calculated according to the formula (36):

$$\sigma = \frac{1}{\sqrt{n\left[\frac{\zeta''(\alpha, x_{min})}{\zeta(\alpha, x_{min})} - \left(\frac{\zeta'(\alpha, x_{min})}{\zeta(\alpha, x_{min})}\right)^2\right]}}, \qquad (S11)$$

where (′) and (″) denote first and second derivatives of function, respectively.

In fact, the maximum likelihood estimation of parameters for other statistical models can be obtained by similar approaches. All maximum likelihood estimations in this work were carried out with the respective code modules from R/Bioconductor library poweRlaw (39).

***2.6.4. Goodness-of-fit test***

Goodness-of-fit test was used for the assessment of plausibility of the statistical hypothesis. This test is based on the measurement of Kolmogorov-Smirnov distance between the distribution of the empirical data and hypothesized model. This distance is compared with the distance measurements for the comparable synthetic data sets drawn from the same model.

Next, the *p*-value is calculated to quantify the plausibility of the hypothesis. In general, the *p*-value is defined as a fraction of synthetic distances that are larger than the empirical distance. If *p* is large ($1 \geq p \geq 0.1$), then the difference between the empirical data and model can be attributed to statistical fluctuations alone; if it is small ($\leq 0.1$), the model is not plausible for the data (36). In this work, the code based on R/Bioconductor library poweRlaw (39) and 1000 bootstrap runs were used for all goodness-of-fit tests.

**2.7. Competitive statistical models**

***2.7.1. Truncated power-law distribution with the exponential cut-off***

The probability mass function for the discrete truncated power-law distribution is

$$p(x) = Cx^{-\alpha}e^{-\lambda x}, \tag{S12}$$

where *C* is the normalization constant, $\alpha$ is the exponent and $\lambda$ is the rate parameter (36).

To normalize this equation the following formula for the constant *C* can be used:

$$C = \frac{e^{\lambda x_{min}}}{\mathrm{lerch}(e^{\lambda}, \alpha, x_{min})}. \tag{S13}$$

***2.7.2. Yule-Simon distribution***

The probability mass function for the discrete Yule-Simon distribution is

$$p(x) = C\frac{\Gamma(x)}{\Gamma(x+\alpha)}, \tag{S14}$$

where *C* is the normalization constant, Γ is the gamma function and $\alpha$ is the exponent (36, 42).

This distribution can be normalized by

$$C = (\alpha - 1)\frac{\Gamma(x_{min} + \alpha - 1)}{\Gamma(x_{min})}. \tag{S15}$$

***2.7.3. Exponential distribution***

The probability mass function for the discrete exponential distribution is

$$p(x) = Ce^{-\lambda x}, \tag{S16}$$

where *C* is the normalization constant and $\lambda$ is the rate parameter (36).

Discrete exponential distribution can be normalized as follows:

$$C = (1 - e^{-\lambda})e^{\lambda x_{min}}. \tag{S17}$$

##### *2.7.4. Stretched exponential distribution (complementary cumulative Weibull distribution)*

The probability density function for the continuous stretched exponential distribution is

$$p(x) = Cx^{\beta-1}e^{-\lambda x^{\beta}}, \quad \text{(S18)}$$

where *C* is the normalization constant, $\lambda$ is the rate parameter and $\beta$ is the stretching exponent (36). This function was discretized and normalized as described by Alstott J. et al. (43).

##### *2.7.5. Log-normal distribution*

The probability density function for the continuous log-normal distribution is

$$p(x) = C\frac{e^{-\frac{(\ln x-\mu)^2}{2\sigma^2}}}{x}, \quad \text{(S19)}$$

where *C* is the normalization constant, $\mu$ is the mean and $\sigma$ is the standard deviation of the natural logarithm of the variable (36).

This function was discretized and normalized as described by Alstott J. et al. (43).

##### *2.7.6. Poisson distribution*

The probability mass function for the discrete Poisson distribution is

$$p(x) = C\frac{\mu^x}{x!}, \quad \text{(S20)}$$

where *C* is the normalization constant and $\mu$ is the mean (36).

To normalize this equation, the following formula for the constant *C* can be used:

$$C = \left[e^{\mu} - \sum_{k=0}^{x_{min}-1}\frac{\mu^k}{k!}\right]^{-1}. \quad \text{(S21)}$$

#### 2.8. Log-likelihood ratio test

Log-likelihood ratio test was used to directly compare alternative statistical models. The likelihood of the data under two competing probability mass functions $p_1(x)$ and $p_2(x)$ was calculated as follows (36, 44):

$$L_1 = \prod_{i=1}^{n} p_1(x_i), \quad \text{(S22)}$$

$$L_2 = \prod_{i=1}^{n} p_2(x_i). \quad \text{(S23)}$$

The likelihood ratio was determined by the equation

$$R = \frac{L_1}{L_2} = \prod_{i=1}^{n}\frac{p_1(x_i)}{p_2(x_i)} \quad \text{(S24)}$$

in its logarithmic form:

$$R = \sum_{i=1}^{n}[lnp_1(x_i) - lnp_2(x_i)] = \sum_{i=1}^{n}\left[l_i^{(1)} - l_i^{(2)}\right], \quad \text{(S25)}$$

where $l_i^{(j)} = lnp_j(x_i)$ is the log-likelihood for a single measurement $x_i$ within distribution *j*.

*R* is positive if the empirical data are more likely in the first statistical model and negative if the second distribution model is better, or zero in the event of a tie.

For non-nested models, statistical significance of *R* was evaluated as proposed (36, 45):

$$p = \frac{1}{\sqrt{2\pi n\sigma^2}}\left[\int_{-\infty}^{-|R|} e^{\frac{-t^2}{2n\sigma^2}}dt + \int_{|R|}^{\infty} e^{\frac{-t^2}{2n\sigma^2}}dt\right] = \left|erfc(\frac{R}{\sqrt{2n\sigma}})\right|, \quad \text{(S26)}$$

where $\sigma$ is given by the equation

$$\sigma^2 = \frac{1}{n}\sum_{i=1}^{n}\left[\left(l_i^{(1)} - l_i^{(2)}\right) - (\bar{l}^{(1)} - \bar{l}^{(2)})\right]^2 \quad \text{(S27)}$$

with

$$\bar{l}^{(1)} = \frac{1}{n}\sum_{i=1}^{n} l_i^{(1)} \quad \text{(S28)}$$

$$\bar{l}^{(2)} = \frac{1}{n}\sum_{i=1}^{n} l_i^{(2)} \quad \text{(S29)}$$

and

$$erfc(z) = 1 - erf(z) = \frac{2}{\sqrt{\pi}}\int_{z}^{\infty} e^{-t^2}dt, \quad \text{(S29)}$$

where *erfc* is the complementary Gaussian error function.

If the *p*-value is < 0.1, *R* is statistically significant. On the other hand, if the *p*-value is > 0.1, the test does not favor one model over the other. As for nested models (for instance, power-law versus truncated power-law), the actual log likelihood ratio was used. In this case, if the p-value is small, then the smaller family can be ruled out. If not, there is no evidence that the larger family is needed to fit the data, although neither can be ruled out (45). All log-likelihood ratio tests in this work were carried out with an in-laboratory developed R code.

**2.9. Information criteria for statistical models**

In some cases, information criteria were used to compare alternative statistical models. First of all, it was Akaike information criterion, or AIC, which can be calculated as follows (46):

$$\mathrm{AIC} = 2\mathrm{k} - 2\ln(\mathrm{L}), \quad \text{(S30)}$$

where *k* is the number of free parameters to be estimated and *L* is the maximized value of the likelihood function of the model.

The second criterion was Schwarz Bayesian information criterion, or BIC, which can be calculated according to a simple equation (47):

$$\mathrm{BIC} = -2\ln(\mathrm{L}) + \mathrm{k}\ln(\mathrm{n}), \quad \text{(S30)}$$

where *L* is the maximized value of the likelihood function of the model, *k* is the number of free parameters to be estimated and *n* is the number of data points in *x*, the number of observations, or equivalently, the sample size.

**2.10. Detection of differential abundance of splicing events by linear modelling**

A primary count matrix of splicing events was prepared for the linear modelling as described in Supplementary Section 2.2. At this step, the count data were logarithmically transformed, the mean-variance relationship was estimated, and the appropriate observational-level weights were calculated. Multiple linear models were fitted to the normalized matrix by least squares with the function *lmFit* from R/Bioconductor library limma (48-49). For fitted models, empirical Bayes statistics of differential abundance of splicing events in two different RNA fractions was calculated with the function *eBayes* from R/Bioconductor library limma (49-50). P-values were adjusted for multiple testing with the method by Benjamini Y. and Hochberg Y., which controls the expected false discovery rate below the specified value (51).

**2.11. Calculation of overlaps between splicing events**

The normalized matrix of identified splicing events was wrapped in a GRanges object (17). Similarly, Cufflinks assembled transcripts and public transcriptional models of human genes were converted in TxFeatures objects (52). These objects were used for the fast extraction of exon-exon junctions in the form of GRanges objects (17). Overlap depth between genomic intervals of different GRanges objects was calculated with the function *findOverlaps* from R/Bioconductor library GenomicRanges (53).

**2.12. Analysis of differential transcript and gene expression**

At the transcript level, we used Cuffdiff FPKM tracking output files (Supplementary Section 2.3) for analysis of differential expression. Herewith, only transcripts with at least 2-fold changes in expression and q-value below 0.1 were annotated as differentially expressed.
At the gene level, genomic coordinates of human exons were retrieved from Ensembl gene models (8). Any overlapping coordinates were collapsed to strand specific coordinates of the group of overlapping exons. The final list of genomic coordinates of a gene-wise individual and/or grouped exons included 328096 items. With this list and RNA-seq data obtained from the mismatch siRNA and anti-RUNX1-RUNX1T1 siRNA treated Kasumi-1 cells, a gene-level read summarization was carried out with a feature Counts software (12). The primary count matrix was prepared for the linear modelling as described in Supplementary Section 2.2 and differential analysis of the gene expression was done in the way described in Supplementary Section 2.10. Output results were parsed and genes with at least 2-fold changes in expression and the q-value below 0.1 were annotated as differentially expressed.

**2.13. Topological analysis of splicing graphs**

In graph theory, a topology is the arrangement of the vertices and edges of a graph (35). A concept of the graph topology can be also applied to splicing graphs to describe the arrangement of splice sites and/or exons during splicing. The topology of splicing graph can be characterized with topological indices. There are two classes of topological indices. Local indices describe the state of each splicing graph element, while global indices characterize a structure of the entire splicing graph.

To describe our splicing graphs, we selected a set of indices that can be interpreted from the biological point of view. Every index was calculated with the respective function from R/Bioconductor library igraph (29) with the exception of complete splicing events (see below).

***2.13.1. Local topological indices***

*Alpha centrality scores*

The alpha centrality is one of generalizations of the eigenvector centrality for directed graphs (55).This index indicates whether there is a tendency for splicing of splice sites (or exons) together with high degrees.

The alpha centrality of the vertices in a graph is defined as the solution of the following matrix equation:

$$x = \alpha A^T x + e, \tag{S31}$$

where $A^T$ is the transpose of adjacency matrix of the graph, *e* is the vector of exogenous sources of status of the exons and $\alpha$ is the relative importance of the endogenous versus exogenous factors.

*Complete events*

A complete event (or a complete alternative splicing event) is a set of all valid paths that form a bubble in a splicing graph (33). A path in a splicing graph is a sequence of adjacent edges between the source vertex *S* and the sink vertex *T* ($S < T$) without traversing any intermediate vertex twice. A valid path between vertices *S* and *T* of a splicing graph is a path followed by at least one experimental transcript. There is a bubble between vertices *S* and *T* of a splicing graph if there are at least two distinct valid paths between vertices *S* and *T* and no other vertices than vertices *S* and *T* are shared by all the valid paths between them. The number of distinct valid paths between vertices *S* and *T* is the dimension of a complete event. We used the function *bubbles* from R/Bioconductor library SplicingGraphs (34) to identify a complete event(-s) in a given gene and to calculate the dimension of each complete event.

*Diversity of vertices*

The diversity *D* of a vertex *i* is defined as the normalized Shannon entropy of the weights of its incident edges. This index can be calculated as follows (29, 56-57):

$$D(i) = \frac{-\sum_{j=1}^{k} p_{ij} log(p_{ij})}{log(k)}, \tag{S32}$$

where *k* is the total degree of vertex *i* and $p_{ij}$ is the proportion of weight $w_{ij}$ of the edge between vertex *i* and vertex *j* in the total weight of the edges of vertex *i*:

$$p_{ij} = \frac{w_{ij}}{\sum_{j=1}^{k} w_{ij}} \tag{S33}$$

We used FPKM values to assign weights to the edges in exon graphs. Alternatively, we weighed the edges with the number of supporting reads in the case of splice site graphs.

*Splicing degrees*

Splicing in-, out, and/or total-degree of each vertex of a splicing graph were calculated as described in Supplementary Section 2.5.

##### *2.13.2. Global topological indices*

*Articulation points*

Articulation points or cut vertices are vertices that increase the number of connected components in a graph after their removal (29). In the context of splicing graphs, an articulation point is usually a constitutive splice site or constitutive exon that connects adjacent complete events and forms a "bottleneck" in the splicing graph.

*Assortativity coefficient*

The assortativity is a preference for vertices of a graph to attach to others that are similar in some way (58). In splicing graph theory, the assortativity coefficient is a measure of preference for splice sites or exons to splice with other vertices that are similar in the value of splicing degrees:

$$r = \frac{\sum_{jk} jk(e_{jk} - q_j^{in} q_k^{out})}{\sigma_{in}\sigma_{out}}, \quad \text{(S34)}$$

where $e_{jk}$ is the probability that a randomly chosen directed edge leads into a vertex of in-degree $j$ and out of a vertex of out-degree $k$, $q^{in}$ is the distribution of the excess in-degree of the vertices that the edges lead into, $q^{out}$ is the distribution of the excess out-degree of the vertices that the edges lead out, $\sigma_{in}$ and $\sigma_{out}$ are standard deviations of distributions $q^{in}$ and $q^{out}$, respectively.

If the assortativity coefficient is 1, the splicing graph is perfectly assortative and splice sites or exons strongly prefer splicing with the similar ones (in terms of values of splicing degrees). When the coefficient is –1, the splicing graph is completely disassortative and splice sites or exons with high degrees are spliced with vertices with low degrees and vice versa. Finally, in the absence of any preference for splicing, the splicing graph is non-assortative and the coefficient is 0.

*Connected components*

The connected component of a graph is a set of pairwise connected vertices (59). Herewith, two vertices are connected if there is a path of edges between them. The connected component is equal to the entire graph or just a sub-graph in strongly and weakly connected graphs, respectively. In the context of splicing graphs, connected components usually represent sub-graphs of individual genes in the exon graph or sub-graphs of interconnected 5' and 3' splice sites in the splice site graph. We transformed our splicing graphs in undirected ones and treated them as weakly connected graphs.

*Graph density*

The density of a splicing graph is the ratio of the number of empirical edges and the number of theoretical edges in a graph (35). Theoretical edges are all possible directed pairwise connections between vertices in a graph. This index can be calculated by a simple equation:

$$D = \frac{E}{V(V-1)}, \quad \text{(S35)}$$

where $E$ is the number of edges and $V$ is the number of vertices in a graph.

*Number of edges*

Number of edges (exons and/or introns depends from type of graph) in a splicing graph.

*Number of vertices*

Number of vertices (splice sites or exons depends from type of graph) in a splicing graph.

*Splicing degree centralization*

The degree centralization of a splicing graph is a measure of how central its most high degree vertex is in relation to how central all other vertices are. The sum of differences in degrees between the most high degree vertex in a graph and all other vertices is calculated in first, then this quantity is divided by the theoretically largest sum of differences in a star graph of the same size. Defined formally, if $D(V_i)$ is a degree of vertex $V_i$, if $D(V_n)$ is the largest degree in the graph, and if

$$max \sum_{i=1}^{V} D(V_n) - D(V_i) \quad \text{(S36)}$$

is the largest sum of differences in vertex degrees for a star graph with the same number of vertices, then the degree centralization of the splicing graph is (60):

$$D_{cen} = \frac{\sum_{i=1}^{V} D(V_n) - D(V_i)}{max \sum_{i=1}^{V} D(V_n) - D(V_i)}. \quad \text{(S37)}$$

The graph with topology resembling a star has the centralization close to 1, whereas a decentralized graph is characterized by having the centralization close to 0. We calculated three variants of the degree centralization index according to the three types of vertex degrees.

##### 2.14. Development of a list of features associated with splice sites

Every splice site was annotated with sequence, sequence-related, functional, and structural features that were extracted from four types of genomic/RNA elements: 100-bp fragment of the upstream exon (USE), 300-bp fragment from the 5' end of the intron (USIF), 300-bp fragment from the 3' end of the intron (DSIF), and 100-bp fragment of the downstream exon (DSE) (see Figure 4A of the main text body of the article). Additionally, each splice site was described with a set of nearest epigenetic markers. In total, the complete list of features included 1680 items.

###### *2.14.1. Sequence features*

*Splice sites scoring*

Genomic coordinates of 5' and 3' splice sites were retrieved from the normalized matrix of experimentally identified splicing events (see Supplementary Section 2.2) and sequences of these sites were extracted from GRCh38/hg38 reference assembly of the Homo sapiens genome with R/Bioconductor library BSgenome.Hsapiens.UCSC.hg38 (61). Position weight matrices for 5' splice sites (9-nucleotide sequence – 3 nucleotides in the exon and 6 nucleotides in the intron) and 3' splice sites (23-nucleotide sequence – 3 nucleotides in the exon and 20 nucleotides in the intron) were downloaded from MIT MaxEnt Splice Site Scoring Server (62). The strength of splice sites was determined by Perl implementation of MaxEntScan algorithms in accordance to the three scoring models: maximum entropy model, first-order Markov model, and weight matrix model. The overall score of exon splice sites was calculated as the sum of individual 5'ss and 3'ss scores averaged over the three models.

*Experimentally verified exonic splicing motifs*

We collected sequences of experimentally verified binding sites that were recognized by DAZAP1 (63-65), ELAVL1 (66-69), FMR1 (70-71), HNRNPA1 (72), HNRNPA2B1 (63-65), HNRNPC (73-75), HNRNPD (76-81), HNRNPL (82-84), HNRNPLL (85-87), HNRNPU (64), SRSF1 (88-91), SRSF2

(88-91), SRSF3 (92-95), SRSF5 (88-91), SRSF6 (88-91), and TARDBP (46-100) splicing related proteins. For each protein and the respective set of sequences, we performed a motif search by the discriminative motif discovery algorithm motifRG (101) against the background set of randomly extracted human intronic and exonic sequences. The primary motif was refined by the function *refinePWMMotif* from R/Bioconductor package motifRG (101) with default settings and was converted into the $\log_2$ position weight matrix with the correction against the background nucleotides frequency. Occurrence of a motif in USE and DSE was determined by the function *countPWM* from R/Bioconductor package Biostrings (102) and was normalized relative to the length of analysed sequences. We used $99^{th}$ quantile of the motif weight distribution as a threshold in the identification of the true motif occurrence.

Additionally, we collected oligomeric sequences that were bound by splicing proteins ELAVL4 (103-105), HNRNPA3 (106), HNRNPC2 (64, 80), HNRNPC1 (80), HNRNPDL (107), HNRNPF (108-110), HNRNPH1 (110-113), HNRNPH2 (110-112, 114), HNRNPH3 (110, 112, 114), HNRNPK (83, 115-116), HNRNPM (117), KHDRBS1 (118-120), KHSRP (121-124), MBNL1 (120, 125-127), PCBP1 (84, 115, 128-129), PCBP2 (83-84, 129-130), QKI (131), RBM25 (132), SF3B1 (133), SFPQ (83, 116, 134-135), SRP54 (136), SRSF4 (137), SRSF7 (92, 138-139), SRSF9 (140-141), SYNCRIP (106, 142), TIA1/TIAL1 (105, 134, 143-147) and YBX1 (65, 148-150). We were not able to calculate position weight matrices of motifs for these proteins due to a limited number of sequences of the experimentally verified binding sites. For this reason, we used an alternative approach in the assessment of the strength of binding sites for the mentioned above splicing proteins, as proposed by Murray J. I. et al. (151):

$$BSS = \frac{\sum_{i=0}^{i=L-k+1} ln(4^k f_{n_i})}{L-k+1}, \tag{S38}$$

where $L$ is the length of the sequence of interest, $k$ is the length of an oligomer (see below) found in the sequence of interest, $f_n$ represents the frequency (within the set of sequences of the experimentally verified binding sites for a given splicing protein) of the oligomer found at the position $i$ in the sequence of interest, and $ln(4^k f_{n_i})$ is a log-odds representation of the degree to which the particular oligomer was enriched within the set of sequences of the experimentally verified binding sites for a given splicing protein. As proposed, we only counted the frequency of all possible pentamers in the sequence of interest and used the frequency of pentamers from the set of sequences of the experimentally verified binding sites for a given splicing protein as the reference (151).

*Bioinformatically predicted exonic splicing motifs*

We selected three different approaches for de novo motifs discovery and identified 25 new motifs that were statistically associated with multi-spliced exons from the Ensembl database. First of all, we used the algorithm GADEM (152) from R/Bioconductor implanted package rGADEM (153). This approach was realized on a subset of exons with splicing degrees equal to or greater than $x_{min}$ from the power-law distribution and with default settings of the software. The second approach was based on the heuristic algorithm *bcrank* from R/Bioconductor package BCRANK (154-155). In this case, short sequences that were overrepresented in ranked exons (in descending order of their splicing degrees) were identified and top motifs were selected for subsequent analysis. Finally, the algorithm *motifRG* from R/Bioconductor implanted in the package motifRG (101) was used with default settings. This algorithm searches for motifs that discriminate the given foreground and background sequences. We used a subset of exons with splicing degrees equal to or greater than $x_{min}$ from the power-law distribution as foreground sequences and other exons from our dataset as background sequences.

All newly identified motifs were converted into $log_2$ position weight matrices with the correction against the background nucleotide frequency. Position weight matrices of Sironi's motifs 1-3 (156) were added to our collection of bioinformatically predicted exonic splicing motifs. Occurrence of these motifs in the sequence of interest was determined as described above. In addition, oligomers with the bioinformatically predicted exonic splicing activity were counted in USE and DSE by the function *vcountPDict* from R/Bioconductor package Biostrings (102) and their frequency was normalized relative to the length of analysed sequences. The list of such oligomers included ESRE hexamers (157), ESRS hexamers (157), ESS decamers (158), PESE octamers (159-160), PESS octamers (159-160), QUEPASA ESEseqs hexamers (161), QUEPASA ESSseqs hexamers (161), and RESCUE ESE hexamers (162).

*Experimentally verified intronic splicing motifs*

We could reconstruct the position weight matrices of motifs for seven splicing proteins that bind intronic sequences: CELF1 (98, 163-165), HNRNPA1 (72), HNRNPA2B1 (63-65), HNRNPL (82-84), SRSF3 (92-95), TARDBP (96-100) and TRA2B (117, 139, 166). We also collected the experimentally verified intronic sequences that are bound by splicing proteins CELF3 (98), ELAVL2 (105), ELAVL4 (103-105), HNRNPA3 (106), HNRNPDL (107), HNRNPF (108-110), HNRNPH1 (110-113), HNRNPH2 (110-112, 114), HNRNPH3 (110, 112, 114), MBNL1 (120, 125, 127), PTBP1 (83, 107, 167-168), PTBP2 (107), QKI (131), RBFOX1 (107, 132, 169-170), RBFOX2 (132, 169-170), RBFOX3 (74, 171), RBM4 (172, 173), SF1 (131, 174), SRSF7 (92, 138-139), SRSF9 (140-141), SYNCRIP (106, 142), TIA1/TIAL1 (105, 134, 143-147), TRA2A (175) and YBX1 (65, 148-150), but for which we could not calculate the position weight matrices because of a limited number of sequences. We used the described above approaches for the determination of occurrence of all these motifs in USIF and DSIF. It should be noted that some splicing proteins did not exhibit any intron/exon preference and bound to both intronic and exonic motifs.

*Bioinformatically predicted intronic splicing motifs*

Oligomers with the bioinformatically predicted splicing activity were counted in USIF and DSIF as described above. The list of such oligomers included Castle's oligomers (176), Culler's ISSs oligomers (177), Das' upstream intronic hexamers (178), Das' downstream intronic hexamers (178), Wang's ISEs hexamers (179), Wang's ISSs decamers (180), Yeo's downstream ISREs (181), and Yeo's upstream ISREs (181).

*Polypyrimidine tract scoring*

For each splicing event, the sequence of the polypyrimidine tract (-30 to -3 positions relative to the acceptor splice site) was extracted from the corresponding DSIF. The strength of U2AF2 binding sites in this sequence was calculated according to Murray J. I. et al. (151):

$$U2AF2_{strength} = \frac{\sum_{i=0}^{i=L-k+1} ln(4^k f_{n_i})}{L-k+1},$$

(S39)

where $L$ is the length of the extracted polypyrimidine tract sequence, $k$ is the length of an oligomer found in the polypyrimidine tract sequence, $f_n$ represents the frequency (within the U2AF2 selected SELEX sequences) of the oligomer found at the position $i$ in the polypyrimidine tract sequence and $ln(4^k f_{n_i})$ is the log-odds representation of the degree to which a particular oligomer was enriched within the U2AF2 selected SELEX sequences (182). The equation S39 is identical to the equation S38, however, values of independent variables in this equation allow us to work only with binding sites for the protein U2AF2. We counted the frequency of all possible pentamers in the polypyrimidine tract and used the frequency of pentamers from the U2AF2 selected SELEX sequences as the reference (151).

Moreover, we collected G- and C-rich 4- to 7-nucleotide sequences overrepresented in intronic regions upstream of the weak polypyrimidine tracts (151). We determined the frequency of these motifs in DSIF (-80 to -30 positions relative to the acceptor splice site) by the function *vcountPDict* from R/Bioconductor package Biostrings (102) and normalized this parameter relative to the length of the analyzed intronic fragment.

*Branchpoint sites scoring*

First of all, we obtained genomic coordinates of 59359 high-confidence human branchpoint sites from work by Mercer T. R. et al. (183). Next, we retrieved the 20-nucleotide sequences surrounding the branchpoints from GRCh38/hg38 reference assembly of the Homo sapiens genome with R/Bioconductor library BSgenome.Hsapiens.UCSC.hg38 (61) and used these sequences as a foreground set in motif discovery analysis. Additionally, we developed a background set of sequences with 10-fold quantitative excess relative to the foreground set. This background set included randomly extracted 100-nucleotide sequences from upstream regions of human introns.

Next, we used the discriminative motif discovery algorithm motifRG with default settings (101) and identified two high confidence motifs associated with human branchpoint sites. These motifs were converted into $log_2$ position weight matrixes with the correction against the background nucleotide frequency and used for the scanning of the sequence of interest. For each splicing event, we extracted a 100-nucleotide sequence (-100 to -1 positions relative to the acceptor splice site) from the corresponding DSIF. For each extracted sequence and each motif, we calculated the maximal and mean motif affinity to the sequence of interest and a number of hits over the score threshold with the function *motifScores* from R/Bioconductor package PWMEnrich (184). As before, we used $99^{th}$ quantile of the motif weight distribution as a threshold in the identification of the true motif occurrence.

*“Short” motifs*

Frequency of $x$-mer (at $x \in [1,4]$) oligonucleotides in the sequence of interest was determined by the function *oligonucleotideFrequency* from R/Bioconductor package Biostrings (102) and normalized relative to the length of the analyzed sequence.

***2.14.2. Sequence-related features***

*Linear density of the minimal free energy of folding*

The sequences of USE, USIF, DSIF, or DSE genomic/RNA elements were extracted from GRCh38/hg38 reference assembly of the *Homo sapiens* genome with R/Bioconductor library BSgenome.Hsapiens.UCSC.hg38 (61). Free energy of folding of these sequences was calculated by the RNAfold tool from ViennaRNA Package (185). Free energy of folding was normalized relative to the length of the analyzed sequence and expressed as a linear density of the minimal free energy (186-187).

*Conservation scores*

BW files with pre-computed conservation scores of the human GRCh38/hg38 reference genome were downloaded via FTP from UCSC Genome Browser (11). We used conservation scores that were calculated with the algorithms phyloP and phastCons after multiz-based multiple alignments of 99 vertebrate genomes to the human genome (188-189). With downloaded BW files, we calculated the minimum, maximum, standard deviation, and mean scores for each sequence of interest.

###### ***2.14.3. Functional features***

We retrieved a functional annotation of human exons from Ensembl gene models (7). Next, we divided all exons into six different classes: non-coding exons (exons that belong to only non-coding transcripts), pure 5’UTR exons, pure CDS exons, pure 3’UTR exons, multitype exons (exons that can be non-coding, 5’UTR, 5’UTR/CDS, CDS, CDS/3’UTR and/or 3’UTR exon depending on transcript), and constitutive/alternative exons. Each splice site was assigned to the class of exon(-s) to which it exactly flanked.

###### ***2.14.4. Structural features***

*Splicing distances*

Splicing distances (length of introns) were directly retrieved from the normalized matrix of the experimentally identified splicing events (see Supplementary Section 2.2).

*Size of exon clusters*

We retrieved genomic coordinates of human exons from Ensembl gene models (7). We grouped exons that overlapped but differed at one or both ends into clusters of exons. The number of exons in a cluster was considered as the size of the cluster. Each splice site was assigned with the size of the exon cluster to which it and its partner splice site(-s) belonged. This metric describes a distribution of constitutive and alternative splice sites along the body of the gene in the immediate vicinity of the splice site of interest.

###### ***2.14.5. Epigenetics features***

In this study, we used data describing the location of five different epigenetics marks in the genome of the siMM or siRR treated Kasumi-1 cells: CpGs islands, DNase I hypersensitivity sites, modified histone H3K9Ac, RNA polymerase II peaks, and RUNX1-RUNX1T1 peaks (see Supplementary Section 1). Distances of the splice sites to the nearest epigenetic marks were measured with the function *distanceToNearest* from R/Bioconductor library GenomicRanges (53).

##### **2.15. Data mining with the random forest meta-classifier**

###### ***2.15.1. Filtration of the primary data matrix***

Our primary data matrix included splicing degrees of splice sites as a dependent response variable and all features described above as an independent predictor of variables. This matrix was filtered against features that had one unique value or features that had both characteristics: i) very few unique values relative to the number of samples and ii) the ratio of the frequency of the most common value to the frequency of the second most common value is large. Highly correlated features were removed with cut-off 0.9 to reduce pair-wise correlations in the matrix. At this step, we used the functionality of R library caret (190).

***2.15.2. Development of a balanced data matrix***

The primary data matrix was big and highly unbalanced because about 94% of splice sites were involved in only one splicing event. In order to balance and decrease the dataset, keeping it representative nevertheless, the following algorithm was used. First, a random sampling of 5000 cases was formed from each class and Euclidian distances between all input vectors in each class were calculated. Next, pairs of cases were selected iteratively ensuring that Euclidian distances between them stayed higher than 0.005 quantile for the distance distribution. Thus, a reasonable number of unsimilar pairs of cases was selected for each class and these cases formed a balanced data matrix.

***2.15.3. Feature importance***

Importance of each feature was determined via calculation of the total decrease in the node impurities from splitting on the feature averaged over all classification trees in random forest. The node impurity was measured with the Gini index. We used R library randomForest in the classification mode at this step (191-192). We also used the balanced data matrix, five independent runs of the random forest meta-classifier, and 1000 classification trees per random forest per run and ranked all features in descending order of importance.

***2.15.4. Feature selection***

We applied a recursive algorithm with five-fold cross-validation to select the minimal required set of important features. We used the function *rfe* from R library caret at this step (190).

***2.15.5. Classification of splice sites***

First of all, the balanced data matrix was reduced to features selected in the previous step. Next, for each run of the random forest meta-classifier, the balanced data matrix was randomly sampled on two sub-matrices: the training set (70% of input matrix) and the test set (30% of input matrix). We used the training set for machine learning, calculation of the proximity matrix and marginal effects of features. The test set was used to assess the classification accuracy. We used R library randomForest in the classification mode at this step (191-192), 1000 trees were grown at each algorithm run and the number of features sampled for splitting up at each node was equal to one third of all features in the input data matrix.

#### 3. SUPPLEMENTARY TABLES

**Supplementary Table S1.** Basic alignment statistics for RNA samples used in this study. The median was chosen as the most appropriate statistical metric to describe a highly asymmetric distribution of fragment lengths.

| Type of RNA | Type of RNA-seq library | RUNX1-RUNX1T1 expression status | RNA sample ID | Reads | | | | | Median of the fragment length |
|---|---|---|---|---|---|---|---|---|---|
| | | | | total | uniquely mapped | support exon-exon junctions | length | mean (± SD) of mapping quality | |
| Nascent RNA | paired-end, stranded | intact | K6_I | 138575558 | 77461425 | 6073937 | 100 | 58.6 ± 0.69 | 145 |
| | | | K6_II | 133934148 | 76280325 | 4956946 | 100 | 58.5 ± 0.86 | 142 |
| | | | K6_III | 131857874 | 69084735 | 5797073 | 100 | 58.6 ± 0.71 | 143 |
| | | down-regulated | K1_I | 116848688 | 71584774 | 4208001 | 100 | 58.7 ± 0.64 | 141 |
| | | | K1_II | 112764036 | 70206194 | 4165294 | 100 | 58.7 ± 0.67 | 152 |
| | | | K1_III | 116515944 | 69809136 | 5002062 | 100 | 58.5 ± 1.18 | 148 |
| Total RNA | paired-end, unstranded | intact | SRR1145838 | 90974676 | 78324713 | 28428577 | 101 | 58.7 ± 0.67 | 194 |
| | | | SRR1145839 | 76614354 | 61994071 | 21600095 | 101 | 58.7 ± 0.67 | 203 |
| | | | SRR1145840 | 91545304 | 78085723 | 27651684 | 101 | 58.8 ± 0.66 | 194 |
| | | down-regulated | SRR1145841 | 76434398 | 65722310 | 22264597 | 101 | 58.7 ± 0.67 | 190 |
| | | | SRR1145842 | 84998500 | 66908910 | 23353412 | 101 | 58.2 ± 0.69 | 199 |
| | | | SRR1145843 | 83931812 | 68971738 | 24378592 | 101 | 58.7 ± 0.67 | 200 |

**Supplementary Table S2.** Maximum values of splicing degrees from different models of human genes.

| Gene models | Basic statistics | | | Splice site level | | Exon level | | |
|---|---|---|---|---|---|---|---|---|
| | #genes | #transcripts | #exons | 5'ss, out-degree | 3'ss, in-degree | total-degree | out-degree | in-degree |
| AceView genes (GRCh37, November 2011) | 72361 | 258927 | 677459 | 15 | 50 | 90 | 52 | 52 |
| ECgene genes (GRCh36.p1, build 1, March 2006) | 66733 | 343405 | 583433 | 12 | 46 | 46 | 12 | 46 |
| Ensembl genes (GRCh38.p7, release 85, July 2016) | 58051 | 198002 | 571400 | 21 | 25 | 85 | 50 | 69 |
| GENCODE genes (GRCh38.p7, release 25, July 2016) | 58037 | 198093 | 571658 | 21 | 25 | 85 | 50 | 69 |
| NCBI RefSeq genes (GRCh38.p7, release 108, June 2016) | 43121 | 170939 | 426977 | 13 | 24 | 33 | 14 | 31 |
| VEGA genes (GRCh38.p8, release 65, June 2016) | 54950 | 202251 | 606215 | 21 | 25 | 84 | 50 | 69 |

**Supplementary Table S3.** Knockdown of the RUNX1-RUNX1T1 fusion gene expression in Kasumi-1 cells did not lead to any change in topology of the whole transcriptome based splice site graphs.

| **Topological index** | **Type of used siRNA** | | **Mann-Whitney U-test, p value** |
|---|---|---|---|
| | **siMM** | **siRR** | |
| **Local indices** | | | |
| Alpha centrality scores | 1.5 ± 0.54 | 1.5 ± 0.54 | 0.9234 |
| Complete events$_{number}$ | 3.9 ± 2.84 | 3.9 ± 2.88 | 0.7422 |
| Complete events$_{dimension}$ | 2.51 ± 0.929 | 2.52 ± 0.935 | 0.2977 |
| Diversity of vertices | 0.663 ± 0.288 | 0.664 ± 0.285 | 0.9306 |
| Splicing degrees$_{total}$ | 1.04 ± 0.216 | 1.04 ± 0.218 | 0.2963 |
| **Global indices** | | | |
| Articulation points | 6467.7 ± 542.3 | 6842.0 ± 785.7 | 0.4 |
| Assortativity coefficient | 0.247 ± 0.002 | 0.245 ± 0.005 | 0.7 |
| Connected components | 92539.3 ± 232.3 | 93011.0 ± 814.3 | 0.7 |
| Graph density | 2.70e-06 ± 7.09e-09 | 2.69e-06± 2.27e-08 | 0.7 |
| Number of edges | 99970.3 ± 889.1 | 100884.7 ± 222.4 | 0.4 |
| Number of vertices | 192420.3 ± 1099.5 | 193805.0 ± 697.2 | 0.2 |
| Splicing degree$_{total}$ centralization | 1.29e-05 ± 8.19e-08 | 1.54e-05 ± 4.32e-08 | 0.1 |

**Supplementary Table S4.** List of features associated with splice sites.

| Class of feature | Feature | 5' ss | 3' ss | USE | USIF | DSIF | DSE |
|---|---|---|---|---|---|---|---|
| 1 | 2 | 3 | 4 | 5 | 6 | 7 | 8 |
| Sequence feature | A | | | ✓ | ✓ | ✓ | ✓ |
| | AA | | | ✓ | ✓ | ✓ | ✓ |
| | AAA | | | ✓ | ✓ | ✓ | ✓ |
| | AAAA | | | ✓ | ✓ | ✓ | ✓ |
| | AAAC | | | ✓ | ✓ | ✓ | ✓ |
| | AAAG | | | ✓ | ✓ | ✓ | ✓ |
| | AAAT | | | ✓ | ✓ | ✓ | ✓ |
| | AAC | | | ✓ | ✓ | ✓ | ✓ |
| | AACA | | | ✓ | ✓ | ✓ | ✓ |
| | AACC | | | ✓ | ✓ | ✓ | ✓ |
| | AACG | | | ✓ | ✓ | ✓ | ✓ |
| | AACT | | | ✓ | ✓ | ✓ | ✓ |
| | AAG | | | ✓ | ✓ | ✓ | ✓ |
| | AAGA | | | ✓ | ✓ | ✓ | ✓ |
| | AAGC | | | ✓ | ✓ | ✓ | ✓ |
| | AAGG | | | ✓ | ✓ | ✓ | ✓ |
| | AAGT | | | ✓ | ✓ | ✓ | ✓ |
| | AAT | | | ✓ | ✓ | ✓ | ✓ |
| | AATA | | | ✓ | ✓ | ✓ | ✓ |
| | AATC | | | ✓ | ✓ | ✓ | ✓ |
| | AATG | | | ✓ | ✓ | ✓ | ✓ |
| | AATT | | | ✓ | ✓ | ✓ | ✓ |
| | AC | | | ✓ | ✓ | ✓ | ✓ |
| | ACA | | | ✓ | ✓ | ✓ | ✓ |
| | ACAA | | | ✓ | ✓ | ✓ | ✓ |
| | ACAC | | | ✓ | ✓ | ✓ | ✓ |
| | ACAG | | | ✓ | ✓ | ✓ | ✓ |
| | ACAT | | | ✓ | ✓ | ✓ | ✓ |
| | ACC | | | ✓ | ✓ | ✓ | ✓ |
| | ACCA | | | ✓ | ✓ | ✓ | ✓ |
| | ACCC | | | ✓ | ✓ | ✓ | ✓ |
| | ACCG | | | ✓ | ✓ | ✓ | ✓ |
| | ACCT | | | ✓ | ✓ | ✓ | ✓ |
| | ACG | | | ✓ | ✓ | ✓ | ✓ |
| | ACGA | | | ✓ | ✓ | ✓ | ✓ |
| | ACGC | | | ✓ | ✓ | ✓ | ✓ |
| | ACGG | | | ✓ | ✓ | ✓ | ✓ |
| | ACGT | | | ✓ | ✓ | ✓ | ✓ |
| | ACT | | | ✓ | ✓ | ✓ | ✓ |
| | ACTA | | | ✓ | ✓ | ✓ | ✓ |
| | ACTC | | | ✓ | ✓ | ✓ | ✓ |
| | ACTG | | | ✓ | ✓ | ✓ | ✓ |
| | ACTT | | | ✓ | ✓ | ✓ | ✓ |
| | AG | | | ✓ | ✓ | ✓ | ✓ |
| | AGA | | | ✓ | ✓ | ✓ | ✓ |

| 1 | 2 | 3 | 4 | 5 | 6 | 7 | 8 |
|---|---|---|---|---|---|---|---|
| | AGAA | | | ✓ | ✓ | ✓ | ✓ |
| | AGAC | | | ✓ | ✓ | ✓ | ✓ |
| | AGAG | | | ✓ | ✓ | ✓ | ✓ |
| | AGAT | | | ✓ | ✓ | ✓ | ✓ |
| | AGC | | | ✓ | ✓ | ✓ | ✓ |
| | AGCA | | | ✓ | ✓ | ✓ | ✓ |
| | AGCC | | | ✓ | ✓ | ✓ | ✓ |
| | AGCG | | | ✓ | ✓ | ✓ | ✓ |
| | AGCT | | | ✓ | ✓ | ✓ | ✓ |
| | AGG | | | ✓ | ✓ | ✓ | ✓ |
| | AGGA | | | ✓ | ✓ | ✓ | ✓ |
| | AGGC | | | ✓ | ✓ | ✓ | ✓ |
| | AGGG | | | ✓ | ✓ | ✓ | ✓ |
| | AGGT | | | ✓ | ✓ | ✓ | ✓ |
| | AGT | | | ✓ | ✓ | ✓ | ✓ |
| | AGTA | | | ✓ | ✓ | ✓ | ✓ |
| | AGTC | | | ✓ | ✓ | ✓ | ✓ |
| | AGTG | | | ✓ | ✓ | ✓ | ✓ |
| | AGTT | | | ✓ | ✓ | ✓ | ✓ |
| | AT | | | ✓ | ✓ | ✓ | ✓ |
| | ATA | | | ✓ | ✓ | ✓ | ✓ |
| | ATAA | | | ✓ | ✓ | ✓ | ✓ |
| | ATAC | | | ✓ | ✓ | ✓ | ✓ |
| | ATAG | | | ✓ | ✓ | ✓ | ✓ |
| | ATAT | | | ✓ | ✓ | ✓ | ✓ |
| | ATC | | | ✓ | ✓ | ✓ | ✓ |
| | ATCA | | | ✓ | ✓ | ✓ | ✓ |
| | ATCC | | | ✓ | ✓ | ✓ | ✓ |
| | ATCG | | | ✓ | ✓ | ✓ | ✓ |
| | ATCT | | | ✓ | ✓ | ✓ | ✓ |
| | ATG | | | ✓ | ✓ | ✓ | ✓ |
| | ATGA | | | ✓ | ✓ | ✓ | ✓ |
| | ATGC | | | ✓ | ✓ | ✓ | ✓ |
| | ATGG | | | ✓ | ✓ | ✓ | ✓ |
| | ATGT | | | ✓ | ✓ | ✓ | ✓ |
| | ATT | | | ✓ | ✓ | ✓ | ✓ |
| | ATTA | | | ✓ | ✓ | ✓ | ✓ |
| | ATTC | | | ✓ | ✓ | ✓ | ✓ |
| | ATTG | | | ✓ | ✓ | ✓ | ✓ |
| | ATTT | | | ✓ | ✓ | ✓ | ✓ |
| | Branchpoint site motif #1, count | | | | | ✓ | |
| | Branchpoint site motif #1, max affinity score | | | | | ✓ | |
| | Branchpoint site motif #1, mean affinity score | | | | | ✓ | |
| | Branchpoint site motif #2, count | | | | | ✓ | |
| | Branchpoint site motif #2, max affinity score | | | | | ✓ | |
| | Branchpoint site motif #2, mean affinity score | | | | | ✓ | |
| | C | | | ✓ | ✓ | ✓ | ✓ |
| | CA | | | ✓ | ✓ | ✓ | ✓ |
| | CAA | | | ✓ | ✓ | ✓ | ✓ |

| 1 | 2 | 3 | 4 | 5 | 6 | 7 | 8 |
|---|---|---|---|---|---|---|---|
| | CAAA | | | ✓ | ✓ | ✓ | ✓ |
| | CAAC | | | ✓ | ✓ | ✓ | ✓ |
| | CAAG | | | ✓ | ✓ | ✓ | ✓ |
| | CAAT | | | ✓ | ✓ | ✓ | ✓ |
| | CAC | | | ✓ | ✓ | ✓ | ✓ |
| | CACA | | | ✓ | ✓ | ✓ | ✓ |
| | CACC | | | ✓ | ✓ | ✓ | ✓ |
| | CACG | | | ✓ | ✓ | ✓ | ✓ |
| | CACT | | | ✓ | ✓ | ✓ | ✓ |
| | CAG | | | ✓ | ✓ | ✓ | ✓ |
| | CAGA | | | ✓ | ✓ | ✓ | ✓ |
| | CAGC | | | ✓ | ✓ | ✓ | ✓ |
| | CAGG | | | ✓ | ✓ | ✓ | ✓ |
| | CAGT | | | ✓ | ✓ | ✓ | ✓ |
| | Castle's oligomers | | | | ✓ | ✓ | |
| | CAT | | | ✓ | ✓ | ✓ | ✓ |
| | CATA | | | ✓ | ✓ | ✓ | ✓ |
| | CATC | | | ✓ | ✓ | ✓ | ✓ |
| | CATG | | | ✓ | ✓ | ✓ | ✓ |
| | CATT | | | ✓ | ✓ | ✓ | ✓ |
| | CC | | | ✓ | ✓ | ✓ | ✓ |
| | CCA | | | ✓ | ✓ | ✓ | ✓ |
| | CCAA | | | ✓ | ✓ | ✓ | ✓ |
| | CCAC | | | ✓ | ✓ | ✓ | ✓ |
| | CCAG | | | ✓ | ✓ | ✓ | ✓ |
| | CCAT | | | ✓ | ✓ | ✓ | ✓ |
| | CCC | | | ✓ | ✓ | ✓ | ✓ |
| | CCCA | | | ✓ | ✓ | ✓ | ✓ |
| | CCCC | | | ✓ | ✓ | ✓ | ✓ |
| | CCCG | | | ✓ | ✓ | ✓ | ✓ |
| | CCCT | | | ✓ | ✓ | ✓ | ✓ |
| | CCG | | | ✓ | ✓ | ✓ | ✓ |
| | CCGA | | | ✓ | ✓ | ✓ | ✓ |
| | CCGC | | | ✓ | ✓ | ✓ | ✓ |
| | CCGG | | | ✓ | ✓ | ✓ | ✓ |
| | CCGT | | | ✓ | ✓ | ✓ | ✓ |
| | CCT | | | ✓ | ✓ | ✓ | ✓ |
| | CCTA | | | ✓ | ✓ | ✓ | ✓ |
| | CCTC | | | ✓ | ✓ | ✓ | ✓ |
| | CCTG | | | ✓ | ✓ | ✓ | ✓ |
| | CCTT | | | ✓ | ✓ | ✓ | ✓ |
| | CELF1 | | | | ✓ | ✓ | |
| | CELF3 | | | | ✓ | ✓ | |
| | CG | | | ✓ | ✓ | ✓ | ✓ |
| | CGA | | | ✓ | ✓ | ✓ | ✓ |
| | CGAA | | | ✓ | ✓ | ✓ | ✓ |
| | CGAC | | | ✓ | ✓ | ✓ | ✓ |
| | CGAG | | | ✓ | ✓ | ✓ | ✓ |
| | CGAT | | | ✓ | ✓ | ✓ | ✓ |

| 1 | 2 | 3 | 4 | 5 | 6 | 7 | 8 |
|---|---|---|---|---|---|---|---|
| | CGC | | | ✓ | ✓ | ✓ | ✓ |
| | CGCA | | | ✓ | ✓ | ✓ | ✓ |
| | CGCC | | | ✓ | ✓ | ✓ | ✓ |
| | CGCG | | | ✓ | ✓ | ✓ | ✓ |
| | CGCT | | | ✓ | ✓ | ✓ | ✓ |
| | CGG | | | ✓ | ✓ | ✓ | ✓ |
| | CGGA | | | ✓ | ✓ | ✓ | ✓ |
| | CGGC | | | ✓ | ✓ | ✓ | ✓ |
| | CGGG | | | ✓ | ✓ | ✓ | ✓ |
| | CGGT | | | ✓ | ✓ | ✓ | ✓ |
| | CGT | | | ✓ | ✓ | ✓ | ✓ |
| | CGTA | | | ✓ | ✓ | ✓ | ✓ |
| | CGTC | | | ✓ | ✓ | ✓ | ✓ |
| | CGTG | | | ✓ | ✓ | ✓ | ✓ |
| | CGTT | | | ✓ | ✓ | ✓ | ✓ |
| | CT | | | ✓ | ✓ | ✓ | ✓ |
| | CTA | | | ✓ | ✓ | ✓ | ✓ |
| | CTAA | | | ✓ | ✓ | ✓ | ✓ |
| | CTAC | | | ✓ | ✓ | ✓ | ✓ |
| | CTAG | | | ✓ | ✓ | ✓ | ✓ |
| | CTAT | | | ✓ | ✓ | ✓ | ✓ |
| | CTC | | | ✓ | ✓ | ✓ | ✓ |
| | CTCA | | | ✓ | ✓ | ✓ | ✓ |
| | CTCC | | | ✓ | ✓ | ✓ | ✓ |
| | CTCG | | | ✓ | ✓ | ✓ | ✓ |
| | CTCT | | | ✓ | ✓ | ✓ | ✓ |
| | CTG | | | ✓ | ✓ | ✓ | ✓ |
| | CTGA | | | ✓ | ✓ | ✓ | ✓ |
| | CTGC | | | ✓ | ✓ | ✓ | ✓ |
| | CTGG | | | ✓ | ✓ | ✓ | ✓ |
| | CTGT | | | ✓ | ✓ | ✓ | ✓ |
| | CTT | | | ✓ | ✓ | ✓ | ✓ |
| | CTTA | | | ✓ | ✓ | ✓ | ✓ |
| | CTTC | | | ✓ | ✓ | ✓ | ✓ |
| | CTTG | | | ✓ | ✓ | ✓ | ✓ |
| | CTTT | | | ✓ | ✓ | ✓ | ✓ |
| | Culler's ISSs oligomers | | | | ✓ | ✓ | |
| | Das's downstream intronic hexamers | | | | ✓ | ✓ | |
| | Das's upstream intronic hexamers | | | | ✓ | ✓ | |
| | DAZAP1 | | | ✓ | | | ✓ |
| | ELAVL1 | | | ✓ | | | ✓ |
| | ELAVL2 | | | | ✓ | ✓ | |
| | ELAVL4 | | | ✓ | ✓ | ✓ | ✓ |
| | Fairbrother's ESE hexamers | | | ✓ | | | ✓ |
| | FMR1 | | | ✓ | | | ✓ |
| | G | | | ✓ | ✓ | ✓ | ✓ |
| | G- and C-rich UPY motifs | | | | | ✓ | |
| | GA | | | ✓ | ✓ | ✓ | ✓ |
| | GAA | | | ✓ | ✓ | ✓ | ✓ |

| 1 | 2 | 3 | 4 | 5 | 6 | 7 | 8 |
|---|---|---|---|---|---|---|---|
| | GAAA | | | ✓ | ✓ | ✓ | ✓ |
| | GAAC | | | ✓ | ✓ | ✓ | ✓ |
| | GAAG | | | ✓ | ✓ | ✓ | ✓ |
| | GAAT | | | ✓ | ✓ | ✓ | ✓ |
| | GAC | | | ✓ | ✓ | ✓ | ✓ |
| | GACA | | | ✓ | ✓ | ✓ | ✓ |
| | GACC | | | ✓ | ✓ | ✓ | ✓ |
| | GACG | | | ✓ | ✓ | ✓ | ✓ |
| | GACT | | | ✓ | ✓ | ✓ | ✓ |
| | GAG | | | ✓ | ✓ | ✓ | ✓ |
| | GAGA | | | ✓ | ✓ | ✓ | ✓ |
| | GAGC | | | ✓ | ✓ | ✓ | ✓ |
| | GAGG | | | ✓ | ✓ | ✓ | ✓ |
| | GAGT | | | ✓ | ✓ | ✓ | ✓ |
| | GAT | | | ✓ | ✓ | ✓ | ✓ |
| | GATA | | | ✓ | ✓ | ✓ | ✓ |
| | GATC | | | ✓ | ✓ | ✓ | ✓ |
| | GATG | | | ✓ | ✓ | ✓ | ✓ |
| | GATT | | | ✓ | ✓ | ✓ | ✓ |
| | GC | | | ✓ | ✓ | ✓ | ✓ |
| | GCA | | | ✓ | ✓ | ✓ | ✓ |
| | GCAA | | | ✓ | ✓ | ✓ | ✓ |
| | GCAC | | | ✓ | ✓ | ✓ | ✓ |
| | GCAG | | | ✓ | ✓ | ✓ | ✓ |
| | GCAT | | | ✓ | ✓ | ✓ | ✓ |
| | GCC | | | ✓ | ✓ | ✓ | ✓ |
| | GCCA | | | ✓ | ✓ | ✓ | ✓ |
| | GCCC | | | ✓ | ✓ | ✓ | ✓ |
| | GCCG | | | ✓ | ✓ | ✓ | ✓ |
| | GCCT | | | ✓ | ✓ | ✓ | ✓ |
| | GCG | | | ✓ | ✓ | ✓ | ✓ |
| | GCGA | | | ✓ | ✓ | ✓ | ✓ |
| | GCGC | | | ✓ | ✓ | ✓ | ✓ |
| | GCGG | | | ✓ | ✓ | ✓ | ✓ |
| | GCGT | | | ✓ | ✓ | ✓ | ✓ |
| | GCT | | | ✓ | ✓ | ✓ | ✓ |
| | GCTA | | | ✓ | ✓ | ✓ | ✓ |
| | GCTC | | | ✓ | ✓ | ✓ | ✓ |
| | GCTG | | | ✓ | ✓ | ✓ | ✓ |
| | GCTT | | | ✓ | ✓ | ✓ | ✓ |
| | GG | | | ✓ | ✓ | ✓ | ✓ |
| | GGA | | | ✓ | ✓ | ✓ | ✓ |
| | GGAA | | | ✓ | ✓ | ✓ | ✓ |
| | GGAC | | | ✓ | ✓ | ✓ | ✓ |
| | GGAG | | | ✓ | ✓ | ✓ | ✓ |
| | GGAT | | | ✓ | ✓ | ✓ | ✓ |
| | GGC | | | ✓ | ✓ | ✓ | ✓ |
| | GGCA | | | ✓ | ✓ | ✓ | ✓ |
| | GGCC | | | ✓ | ✓ | ✓ | ✓ |

| 1 | 2 | 3 | 4 | 5 | 6 | 7 | 8 |
|---|---|---|---|---|---|---|---|
| | GGCG | | | ✓ | ✓ | ✓ | ✓ |
| | GGCT | | | ✓ | ✓ | ✓ | ✓ |
| | GGG | | | ✓ | ✓ | ✓ | ✓ |
| | GGGA | | | ✓ | ✓ | ✓ | ✓ |
| | GGGC | | | ✓ | ✓ | ✓ | ✓ |
| | GGGG | | | ✓ | ✓ | ✓ | ✓ |
| | GGGT | | | ✓ | ✓ | ✓ | ✓ |
| | GGT | | | ✓ | ✓ | ✓ | ✓ |
| | GGTA | | | ✓ | ✓ | ✓ | ✓ |
| | GGTC | | | ✓ | ✓ | ✓ | ✓ |
| | GGTG | | | ✓ | ✓ | ✓ | ✓ |
| | GGTT | | | ✓ | ✓ | ✓ | ✓ |
| | Goren's ESRE hexamers | | | ✓ | | | ✓ |
| | Goren's ESRS hexamers | | | ✓ | | | ✓ |
| | GT | | | ✓ | ✓ | ✓ | ✓ |
| | GTA | | | ✓ | ✓ | ✓ | ✓ |
| | GTAA | | | ✓ | ✓ | ✓ | ✓ |
| | GTAC | | | ✓ | ✓ | ✓ | ✓ |
| | GTAG | | | ✓ | ✓ | ✓ | ✓ |
| | GTAT | | | ✓ | ✓ | ✓ | ✓ |
| | GTC | | | ✓ | ✓ | ✓ | ✓ |
| | GTCA | | | ✓ | ✓ | ✓ | ✓ |
| | GTCC | | | ✓ | ✓ | ✓ | ✓ |
| | GTCG | | | ✓ | ✓ | ✓ | ✓ |
| | GTCT | | | ✓ | ✓ | ✓ | ✓ |
| | GTG | | | ✓ | ✓ | ✓ | ✓ |
| | GTGA | | | ✓ | ✓ | ✓ | ✓ |
| | GTGC | | | ✓ | ✓ | ✓ | ✓ |
| | GTGG | | | ✓ | ✓ | ✓ | ✓ |
| | GTGT | | | ✓ | ✓ | ✓ | ✓ |
| | GTT | | | ✓ | ✓ | ✓ | ✓ |
| | GTTA | | | ✓ | ✓ | ✓ | ✓ |
| | GTTC | | | ✓ | ✓ | ✓ | ✓ |
| | GTTG | | | ✓ | ✓ | ✓ | ✓ |
| | GTTT | | | ✓ | ✓ | ✓ | ✓ |
| | highECI motif #1 | | | ✓ | | | ✓ |
| | highECI motif #10 | | | ✓ | | | ✓ |
| | highECI motif #11 | | | ✓ | | | ✓ |
| | highECI motif #12 | | | ✓ | | | ✓ |
| | highECI motif #13 | | | ✓ | | | ✓ |
| | highECI motif #14 | | | ✓ | | | ✓ |
| | highECI motif #15 | | | ✓ | | | ✓ |
| | highECI motif #16 | | | ✓ | | | ✓ |
| | highECI motif #17 | | | ✓ | | | ✓ |
| | highECI motif #18 | | | ✓ | | | ✓ |
| | highECI motif #19 | | | ✓ | | | ✓ |
| | highECI motif #2 | | | ✓ | | | ✓ |
| | highECI motif #20 | | | ✓ | | | ✓ |
| | highECI motif #21 | | | ✓ | | | ✓ |

| 1 | 2 | 3 | 4 | 5 | 6 | 7 | 8 |
|---|---|---|---|---|---|---|---|
| | highECI motif #22 | | | ✓ | | | ✓ |
| | highECI motif #23 | | | ✓ | | | ✓ |
| | highECI motif #24 | | | ✓ | | | ✓ |
| | highECI motif #25 | | | ✓ | | | ✓ |
| | highECI motif #3 | | | ✓ | | | ✓ |
| | highECI motif #4 | | | ✓ | | | ✓ |
| | highECI motif #5 | | | ✓ | | | ✓ |
| | highECI motif #6 | | | ✓ | | | ✓ |
| | highECI motif #7 | | | ✓ | | | ✓ |
| | highECI motif #8 | | | ✓ | | | ✓ |
| | highECI motif #9 | | | ✓ | | | ✓ |
| | HNRNPA1 | | | ✓ | ✓ | ✓ | ✓ |
| | HNRNPA2B1 | | | ✓ | ✓ | ✓ | ✓ |
| | HNRNPA3 | | | ✓ | ✓ | ✓ | ✓ |
| | HNRNPC | | | ✓ | | | ✓ |
| | HNRNPC1 | | | ✓ | | | ✓ |
| | HNRNPC2 | | | ✓ | | | ✓ |
| | HNRNPD | | | ✓ | | | ✓ |
| | HNRNPDL | | | ✓ | ✓ | ✓ | ✓ |
| | HNRNPF | | | ✓ | ✓ | ✓ | ✓ |
| | HNRNPH1 | | | ✓ | ✓ | ✓ | ✓ |
| | HNRNPH2 | | | ✓ | ✓ | ✓ | ✓ |
| | HNRNPH3 | | | ✓ | ✓ | ✓ | ✓ |
| | HNRNPK | | | ✓ | | | ✓ |
| | HNRNPL | | | ✓ | ✓ | ✓ | ✓ |
| | HNRNPLL | | | ✓ | | | ✓ |
| | HNRNPM | | | ✓ | | | ✓ |
| | HNRNPU | | | ✓ | | | ✓ |
| | KHDRBS1 | | | ✓ | | | ✓ |
| | KHSRP | | | ✓ | | | ✓ |
| | MBNL1 | | | ✓ | ✓ | ✓ | ✓ |
| | PCBP1 | | | ✓ | | | ✓ |
| | PCBP2 | | | ✓ | | | ✓ |
| | PTBP1 | | | ✓ | ✓ | ✓ | ✓ |
| | PTBP2 | | | | ✓ | ✓ | |
| | QKI | | | ✓ | ✓ | ✓ | ✓ |
| | RBFOX1 | | | | ✓ | ✓ | |
| | RBFOX2 | | | | ✓ | ✓ | |
| | RBFOX3 | | | | ✓ | ✓ | |
| | RBM25 | | | ✓ | | | ✓ |
| | RBM4 | | | | ✓ | ✓ | |
| | SF1 | | | | ✓ | ✓ | |
| | SF3B2 | | | ✓ | | | ✓ |
| | SFPQ | | | ✓ | | | ✓ |
| | Shengdong's ESEseqs hexamers | | | ✓ | | | ✓ |
| | Shengdong's ESSseqs hexamers | | | ✓ | | | ✓ |
| | Sironi's motif #1 | | | ✓ | | | ✓ |
| | Sironi's motif #2 | | | ✓ | | | ✓ |
| | Sironi's motif #3 | | | ✓ | | | ✓ |

| 1 | 2 | 3 | 4 | 5 | 6 | 7 | 8 |
|---|---|---|---|---|---|---|---|
| | Splice site score | ✓ | ✓ | | | | |
| | SRP54 | | | ✓ | | | ✓ |
| | SRSF1 | | | ✓ | | | ✓ |
| | SRSF2 | | | ✓ | | | ✓ |
| | SRSF3 | | | ✓ | ✓ | ✓ | ✓ |
| | SRSF4 | | | ✓ | | | ✓ |
| | SRSF5 | | | ✓ | | | ✓ |
| | SRSF6 | | | ✓ | | | ✓ |
| | SRSF7 | | | ✓ | ✓ | ✓ | ✓ |
| | SRSF9 | | | ✓ | ✓ | ✓ | ✓ |
| | SYNCRIP | | | ✓ | ✓ | ✓ | ✓ |
| | T | | | ✓ | ✓ | ✓ | ✓ |
| | TA | | | ✓ | ✓ | ✓ | ✓ |
| | TAA | | | ✓ | ✓ | ✓ | ✓ |
| | TAAA | | | ✓ | ✓ | ✓ | ✓ |
| | TAAC | | | ✓ | ✓ | ✓ | ✓ |
| | TAAG | | | ✓ | ✓ | ✓ | ✓ |
| | TAAT | | | ✓ | ✓ | ✓ | ✓ |
| | TAC | | | ✓ | ✓ | ✓ | ✓ |
| | TACA | | | ✓ | ✓ | ✓ | ✓ |
| | TACC | | | ✓ | ✓ | ✓ | ✓ |
| | TACG | | | ✓ | ✓ | ✓ | ✓ |
| | TACT | | | ✓ | ✓ | ✓ | ✓ |
| | TAG | | | ✓ | ✓ | ✓ | ✓ |
| | TAGA | | | ✓ | ✓ | ✓ | ✓ |
| | TAGC | | | ✓ | ✓ | ✓ | ✓ |
| | TAGG | | | ✓ | ✓ | ✓ | ✓ |
| | TAGT | | | ✓ | ✓ | ✓ | ✓ |
| | TARDBP | | | ✓ | ✓ | ✓ | ✓ |
| | TAT | | | ✓ | ✓ | ✓ | ✓ |
| | TATA | | | ✓ | ✓ | ✓ | ✓ |
| | TATC | | | ✓ | ✓ | ✓ | ✓ |
| | TATG | | | ✓ | ✓ | ✓ | ✓ |
| | TATT | | | ✓ | ✓ | ✓ | ✓ |
| | TC | | | ✓ | ✓ | ✓ | ✓ |
| | TCA | | | ✓ | ✓ | ✓ | ✓ |
| | TCAA | | | ✓ | ✓ | ✓ | ✓ |
| | TCAC | | | ✓ | ✓ | ✓ | ✓ |
| | TCAG | | | ✓ | ✓ | ✓ | ✓ |
| | TCAT | | | ✓ | ✓ | ✓ | ✓ |
| | TCC | | | ✓ | ✓ | ✓ | ✓ |
| | TCCA | | | ✓ | ✓ | ✓ | ✓ |
| | TCCC | | | ✓ | ✓ | ✓ | ✓ |
| | TCCG | | | ✓ | ✓ | ✓ | ✓ |
| | TCCT | | | ✓ | ✓ | ✓ | ✓ |
| | TCG | | | ✓ | ✓ | ✓ | ✓ |
| | TCGA | | | ✓ | ✓ | ✓ | ✓ |
| | TCGC | | | ✓ | ✓ | ✓ | ✓ |
| | TCGG | | | ✓ | ✓ | ✓ | ✓ |

| 1 | 2 | 3 | 4 | 5 | 6 | 7 | 8 |
|---|---|---|---|---|---|---|---|
| | TCGT | | | ✓ | ✓ | ✓ | ✓ |
| | TCT | | | ✓ | ✓ | ✓ | ✓ |
| | TCTA | | | ✓ | ✓ | ✓ | ✓ |
| | TCTC | | | ✓ | ✓ | ✓ | ✓ |
| | TCTG | | | ✓ | ✓ | ✓ | ✓ |
| | TCTT | | | ✓ | ✓ | ✓ | ✓ |
| | TG | | | ✓ | ✓ | ✓ | ✓ |
| | TGA | | | ✓ | ✓ | ✓ | ✓ |
| | TGAA | | | ✓ | ✓ | ✓ | ✓ |
| | TGAC | | | ✓ | ✓ | ✓ | ✓ |
| | TGAG | | | ✓ | ✓ | ✓ | ✓ |
| | TGAT | | | ✓ | ✓ | ✓ | ✓ |
| | TGC | | | ✓ | ✓ | ✓ | ✓ |
| | TGCA | | | ✓ | ✓ | ✓ | ✓ |
| | TGCC | | | ✓ | ✓ | ✓ | ✓ |
| | TGCG | | | ✓ | ✓ | ✓ | ✓ |
| | TGCT | | | ✓ | ✓ | ✓ | ✓ |
| | TGG | | | ✓ | ✓ | ✓ | ✓ |
| | TGGA | | | ✓ | ✓ | ✓ | ✓ |
| | TGGC | | | ✓ | ✓ | ✓ | ✓ |
| | TGGG | | | ✓ | ✓ | ✓ | ✓ |
| | TGGT | | | ✓ | ✓ | ✓ | ✓ |
| | TGT | | | ✓ | ✓ | ✓ | ✓ |
| | TGTA | | | ✓ | ✓ | ✓ | ✓ |
| | TGTC | | | ✓ | ✓ | ✓ | ✓ |
| | TGTG | | | ✓ | ✓ | ✓ | ✓ |
| | TGTT | | | ✓ | ✓ | ✓ | ✓ |
| | TIA1 | | | ✓ | ✓ | ✓ | ✓ |
| | TIAL1 | | | ✓ | ✓ | ✓ | ✓ |
| | TRA2B | | | | ✓ | ✓ | |
| | TT | | | ✓ | ✓ | ✓ | ✓ |
| | TTA | | | ✓ | ✓ | ✓ | ✓ |
| | TTAA | | | ✓ | ✓ | ✓ | ✓ |
| | TTAC | | | ✓ | ✓ | ✓ | ✓ |
| | TTAG | | | ✓ | ✓ | ✓ | ✓ |
| | TTAT | | | ✓ | ✓ | ✓ | ✓ |
| | TTC | | | ✓ | ✓ | ✓ | ✓ |
| | TTCA | | | ✓ | ✓ | ✓ | ✓ |
| | TTCC | | | ✓ | ✓ | ✓ | ✓ |
| | TTCG | | | ✓ | ✓ | ✓ | ✓ |
| | TTCT | | | ✓ | ✓ | ✓ | ✓ |
| | TTG | | | ✓ | ✓ | ✓ | ✓ |
| | TTGA | | | ✓ | ✓ | ✓ | ✓ |
| | TTGC | | | ✓ | ✓ | ✓ | ✓ |
| | TTGG | | | ✓ | ✓ | ✓ | ✓ |
| | TTGT | | | ✓ | ✓ | ✓ | ✓ |
| | TTT | | | ✓ | ✓ | ✓ | ✓ |
| | TTTA | | | ✓ | ✓ | ✓ | ✓ |
| | TTTC | | | ✓ | ✓ | ✓ | ✓ |

| 1 | 2 | 3 | 4 | 5 | 6 | 7 | 8 |
|---|---|---|---|---|---|---|---|
| | TTTG | | | ✓ | ✓ | ✓ | ✓ |
| | TTTT | | | ✓ | ✓ | ✓ | ✓ |
| | U2AF2 | | | | | ✓ | |
| | Wang's ESS decamers | | | ✓ | | | ✓ |
| | Wang's ISEs hexamers | | | | ✓ | ✓ | |
| | Wang's ISSs decamers | | | | ✓ | ✓ | |
| | YBX1 | | | ✓ | ✓ | ✓ | ✓ |
| | Yeo's downstream ISREs | | | | ✓ | ✓ | |
| | Yeo's upstream ISREs | | | | ✓ | ✓ | |
| | Zhang's PESE octamers | | | ✓ | | | ✓ |
| | Zhang's PESS octamers | | | ✓ | | | ✓ |
| Sequence related feature | Linear density of the minimal free energy | | | ✓ | ✓ | ✓ | ✓ |
| | phastCons score, min | | | ✓ | ✓ | ✓ | ✓ |
| | phastCons score, max | | | ✓ | ✓ | ✓ | ✓ |
| | phastCons score, mean | | | ✓ | ✓ | ✓ | ✓ |
| | phastCons score, SD | | | ✓ | ✓ | ✓ | ✓ |
| | phyloP score, min | | | ✓ | ✓ | ✓ | ✓ |
| | phyloP score, max | | | ✓ | ✓ | ✓ | ✓ |
| | phyloP score, mean | | | ✓ | ✓ | ✓ | ✓ |
| | phyloP score, SD | | | ✓ | ✓ | ✓ | ✓ |
| Functional feature | 5’UTR | | | ✓ | | | ✓ |
| | CDS | | | ✓ | | | ✓ |
| | 3’UTR | | | ✓ | | | ✓ |
| | Multitype | | | ✓ | | | ✓ |
| | Non-coding | | | ✓ | | | ✓ |
| | Constitutive | | | ✓ | | | ✓ |
| Structural feature | Splicing distance | ✓ | ✓ | | | | |
| | Size of exons cluster | ✓ | ✓ | | | | |
| Epigenetic feature | Distance to CpGs island | ✓ | ✓ | | | | |
| | Distance to DNase I hypersensitivity site | ✓ | ✓ | | | | |
| | Distance to H3K9Ac | ✓ | ✓ | | | | |
| | Distance to RNA polymerase II peak | ✓ | ✓ | | | | |
| | Distance to RUNX1-RUNX1T1 peak | ✓ | ✓ | | | | |

**Supplementary Table S5.** Classification accuracy of splice sites from different datasets with the random forest meta-classifier.

| Treatment conditions | Dataset | Classification accuracy, mean ± SD (from 5 independent runs of the meta-classifier) | | |
|---|---|---|---|---|
| | | **overall** | **unisplice** | **multisplice** |
| **siMM** | **5' splice sites** | | | |
| | Original data | 0.799 ± 0.002 | 0.824 ± 0.009 | 0.775 ± 0.006 |
| | SC.1 + SC.2 | 0.999 ± 0.001 | 0.999 ± 0.001 | 1.000 ± 0.000 |
| | MC.1 + MC.2 | 0.892 ± 0.008 | 0.912 ± 0.009 | 0.873 ± 0.010 |
| | SC as train versus MC as test | 0.140 ± 0.006 | 0.130 ± 0.014 | 0.151 ± 0.003 |
| | MC as train versus SC as test | 0.016 ± 0.003 | 0.020 ± 0.006 | 0.011 ± 0.001 |
| | **3' splice sites** | | | |
| | Original data | 0.802 ± 0.003 | 0.802 ± 0.006 | 0.801 ± 0.006 |
| | SC.1 + SC.2 | 0.999 ± 0.001 | 0.999 ± 0.001 | 1.000 ± 0.000 |
| | MC.1 + MC.2 | 0.911 ± 0.007 | 0.923 ± 0.007 | 0.899 ± 0.011 |
| | SC as train versus MC as test | 0.093 ± 0.005 | 0.080 ± 0.011 | 0.105 ± 0.002 |
| | MC as train versus SC as test | 0.005 ± 0.001 | 0.008 ± 0.001 | 0.001 ± 0.001 |
| **siRR** | **5' splice sites** | | | |
| | Original data | 0.798 ± 0.007 | 0.820 ± 0.003 | 0.776 ± 0.013 |
| | SC.1 + SC.2 | 0.999 ± 0.001 | 0.998 ± 0.001 | 1.000 ± 0.000 |
| | MC.1 + MC.2 | 0.893 ± 0.005 | 0.909 ± 0.009 | 0.877 ± 0.010 |
| | SC as train versus MC as test | 0.129 ± 0.002 | 0.084 ± 0.006 | 0.174 ± 0.003 |
| | MC as train versus SC as test | 0.014 ± 0.003 | 0.013 ± 0.003 | 0.014 ± 0.003 |
| | **3' splice sites** | | | |
| | Original data | 0.797 ± 0.013 | 0.788 ± 0.021 | 0.806 ± 0.008 |
| | SC.1 + SC.2 | 0.999 ± 0.001 | 0.998 ± 0.002 | 1.000 ± 0.000 |
| | MC.1 + MC.2 | 0.908 ± 0.005 | 0.927 ± 0.004 | 0.890 ± 0.010 |
| | SC as train versus MC as test | 0.100 ± 0.002 | 0.094 ± 0.005 | 0.106 ± 0.001 |
| | MC as train versus SC as test | 0.005 ± 0.001 | 0.008 ± 0.002 | 0.001 ± 0.000 |

#### 4. SUPPLEMENTARY FIGURES

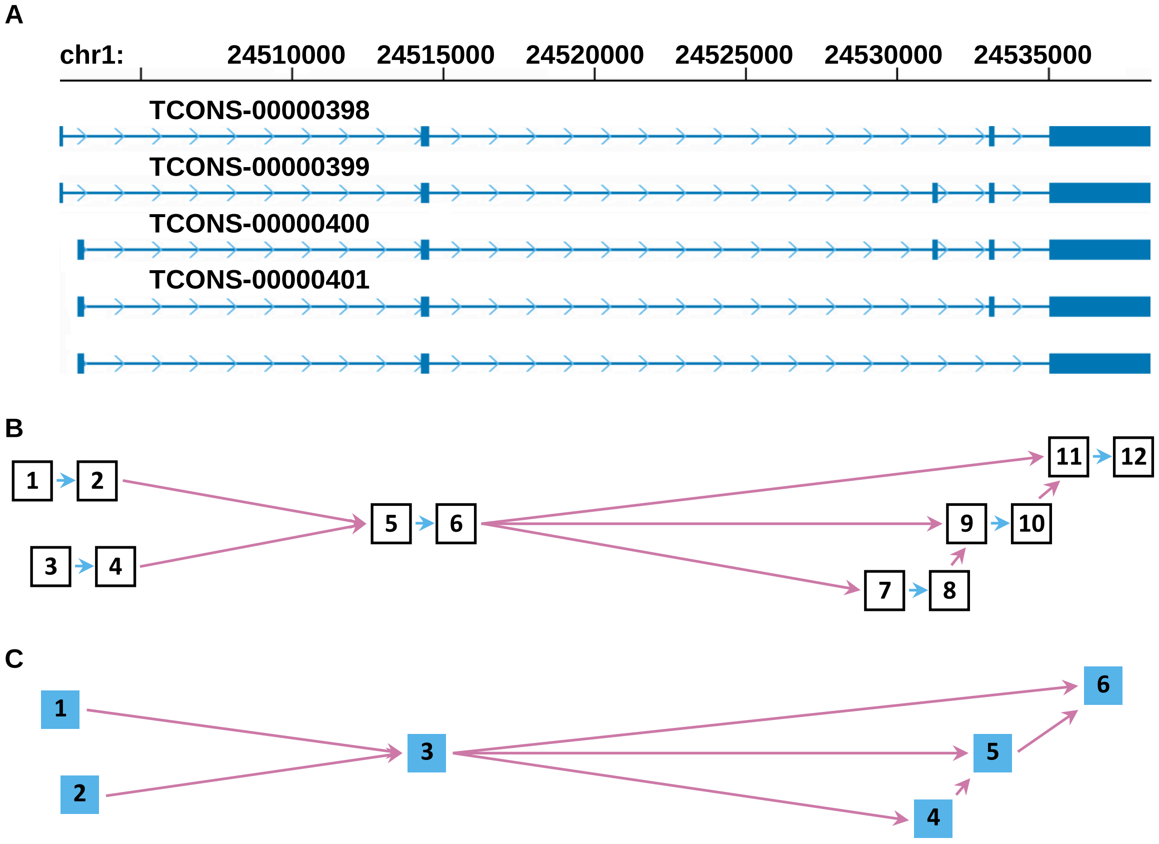


**Supplementary Figure S1.** Splicing graphs. (A) A graphical representation of the traditional (linear) transcriptional model of the human gene organisation. For this figure, the RCAN3 gene was used as an example. Genomic coordinates of the gene are indicated on the top of the figure. Exons are shown with scaled blue rectangles, horizontal scaled blue lines represent introns, direction of transcription is shown by sky blue arrows. Transcripts were assembled with Cufflinks from RNA-seq reads, and original Cufflinks IDs of transcripts were used. Reads were obtained from the mismatch siRNA treated Kasumi-1 cells. (B) Compact representation of the diversity of Cufflinks-based transcripts of the RCAN3 gene with the splice site graph. In this graph, each splice site is a vertex of graph shown with a numbered white rectangle. Intermediate exons and introns are the edges of the graph. They are depicted with sky blue and reddish purple arrows, respectively. From this graph, splicing degree of each splice site can be calculated. For instance, 3’ splice site #5 has in-degree = 2 (two ingoing adjacent edges 2→5 and 4→5) and 5’ splice site #6 has out-degree = 3 (three outgoing adjacent edges 6→7, 6→9 and 6→11). (C) An exon graph-based representation of the same Cufflinks assembled transcripts of the RCAN3 gene. In this case, each exon is a vertex of graph shown with a numbered sky blue rectangle. The connecting edges of this graph are introns. They are depicted with reddish purple arrows. Again, splicing degree of each vertex can be directly calculated from the structure of the graph. For example, usage of the “bottleneck” exon #3 in splicing can be described with three indices: in-degree = 2, out-degree = 3 and total-degree = 5.

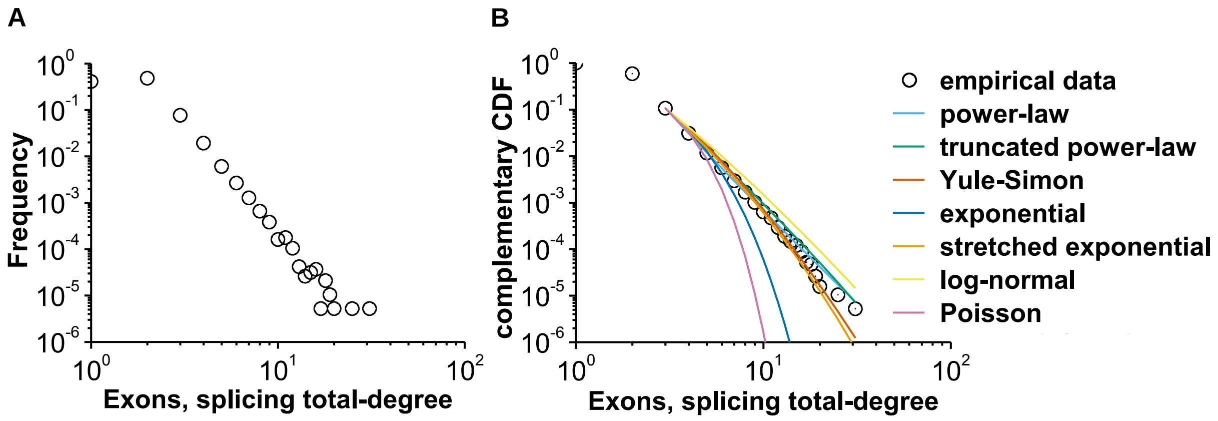


**Supplementary Figure S2.** Usage of exons in alternative splicing follows the power-law in the human transcriptome. (A) Frequency plot of exons with different values of splicing total-degree. This plot is based on the whole dataset of human transcripts from GenBank. (B) Log-log plot of the complementary cumulative distribution function, or complementary CDF, for the splicing total-degrees of exons from the whole dataset of human GenBank transcripts. In this figure, the best fits of competing statistical models are shown. The power-law is the most plausible model to describe heavy right tail of the empirical data.

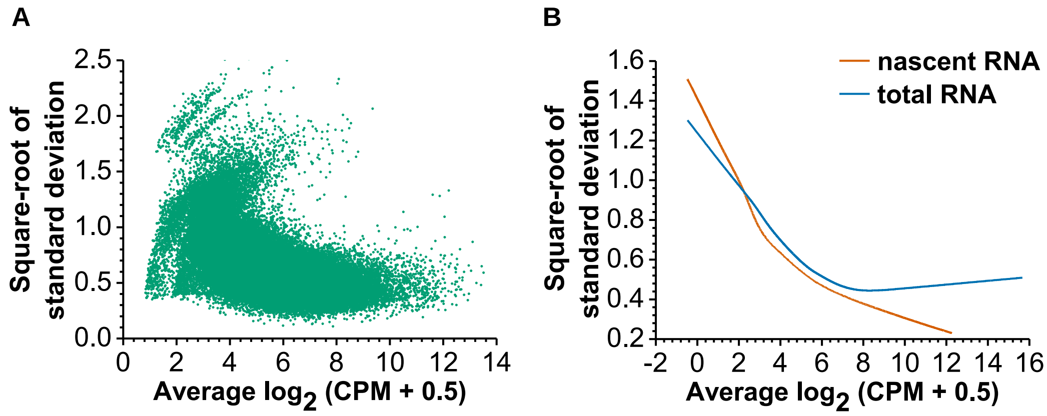


**Supplementary Figure S3.** Mean-variance relationships for nascent RNA and total RNA datasets. (A) Scatterplot of the mean-variance relationship. This plot is based on pooled together nascent and total RNAs dataset. Splicing event-wise square-roots of standard deviations were plotted against the average $log_2$-transformed CPM. (B) Nascent RNA and total RNA datasets differ in the level of technical and biological variability. Technical variability is present at low values of CPM, while the biological variability is visible at high CPMs. It can be demonstrated with the robust lowess (locally weighted regression) voom trend that fits to points of the mean-variance relationship. The nascent RNA dataset is characterized by a moderate degree of the technical variability and a very low biological variation. In contrast, the total RNA dataset shows a low level of technical variability and a high-level biological variation.

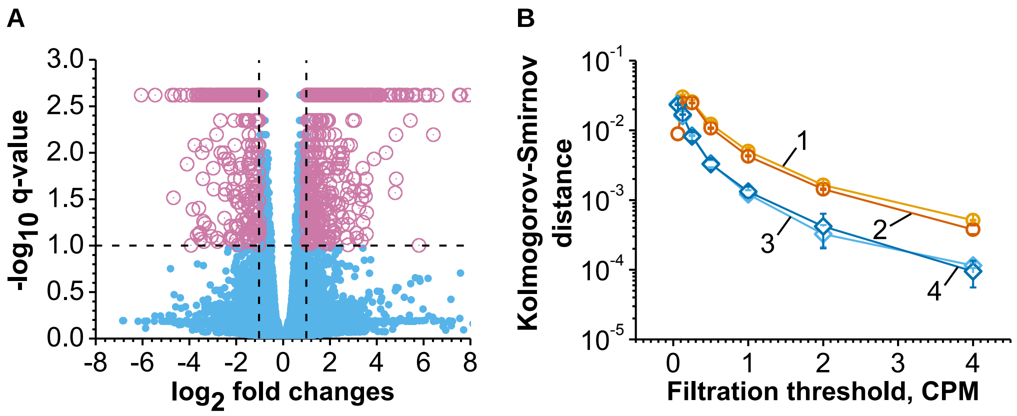


**Supplementary Figure S4.** Large-scale changes in the transcript expression did not change the shape of the distribution of splice sites degrees in the Kasumi-1 transcriptome. (A) At the level of individual transcripts, there is a significant difference between expression of 5995 (of 44638) Cufflinks assembled transcripts in the siMM and siRR treated leukemia cells. The horizontal dotted line corresponds to the q-value = 0.1; vertical dotted lines mark the 2-fold change in expression. (B) The power-law model can be fitted equally well to datasets obtained from both the siMM and siRR treated Kasumi-1 cells. In this plot, empirical distributions of 3' splice sites in-degrees were objects of analysis. This plot is based on three independent sequencing results of the transcriptome of leukemia cells. Each data point is shown as the mean ±SD. Lines 1 and 2 correspond to the nascent RNA dataset and lines 3 and 4 represent the total RNA dataset. Additionally, lines 1 and 3 show the result from the siMM treated Kasumi-1 cells and lines 2 and 4 correspond to data from the siRR treated Kasumi-1 cells.

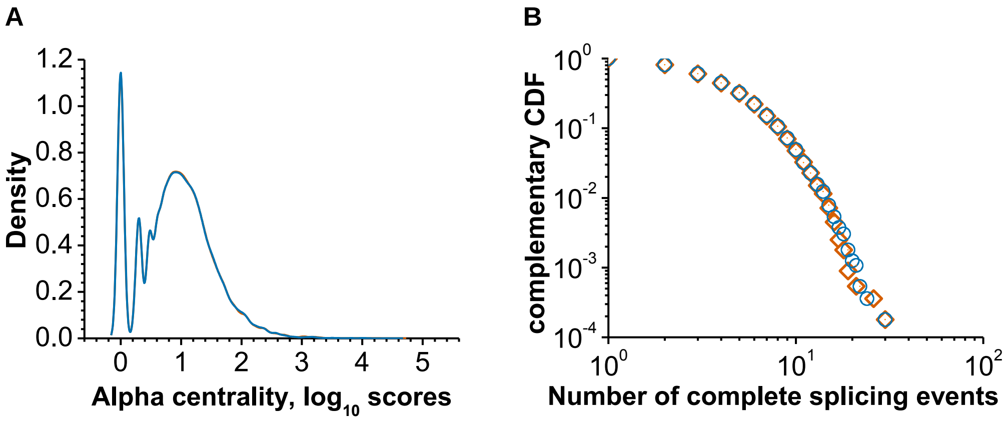


**Supplementary Figure S5.** siRNA-mediated knockdown of the RUNX1-RUNX1T1 fusion gene expression in Kasumi-1 cells did not change distribution of local topological indices of the whole transcriptome based splicing graphs. (A) Distributions of the $log_{10}$-transformed alpha centrality scores for two exon graphs. The alpha centrality scores were calculated as described in Supplementary Section 2.13. Exon graphs were reconstructed from Cufflinks assembled transcriptomes of the mismatch (—) and anti-RUNX1-RUNX1T1 (—) siRNAs treated Kasumi-1 cells. In this picture, distribution lines almost completely overlap. (A) Log-log plot of the complementary CDF for the splice site subgraph-wise number of complete splicing events. Complete splicing events were identified in splice site graphs as described in Supplementary Section 2.13. Splice site graphs were inferred from transcriptomes of the siMM (◇) and siRR (●) treated leukemia cells.

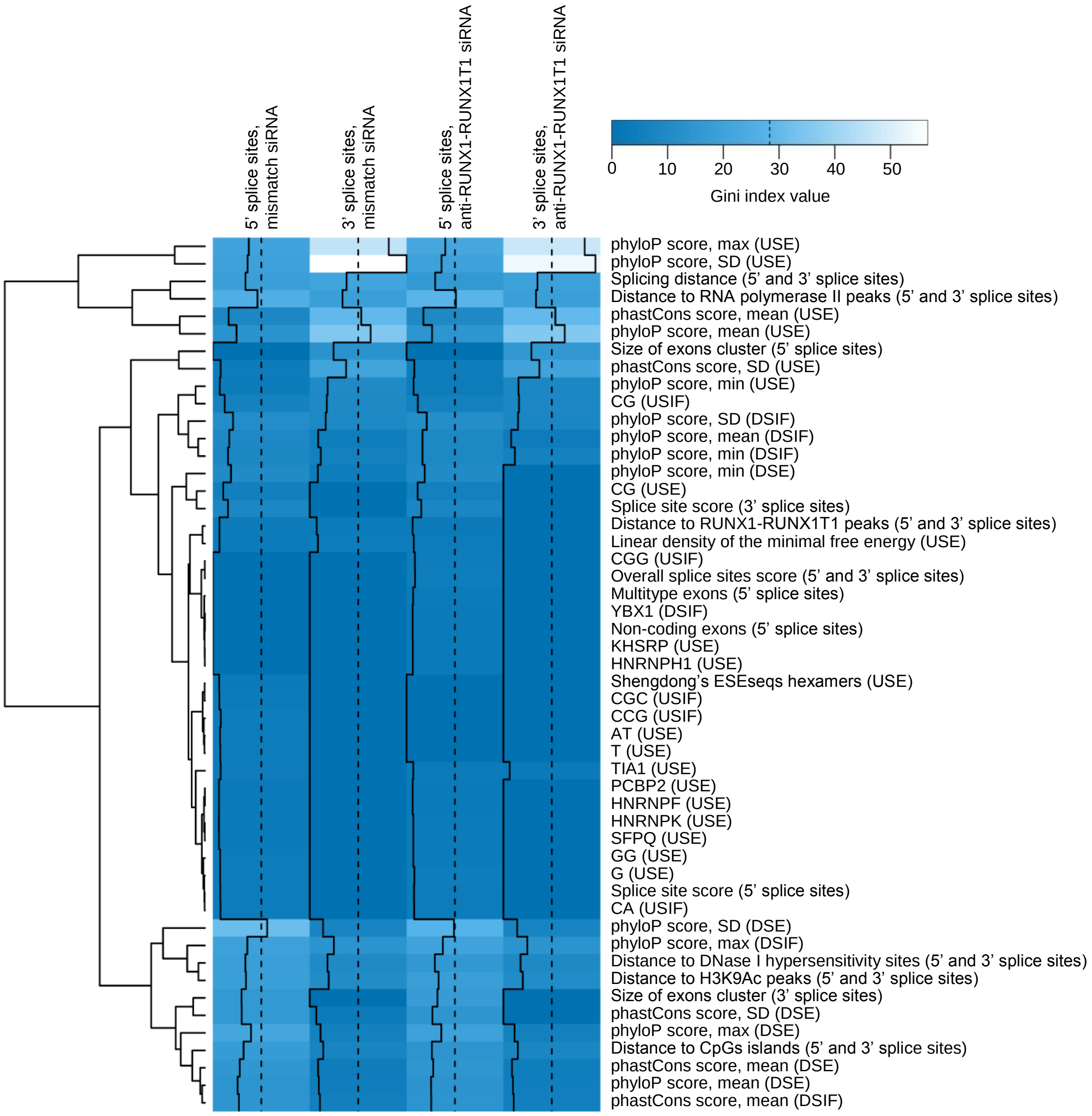


**Supplementary Figure S6.** All selected features of splice sites can be assigned into three clusters. In parentheses, genetic elements are given. Distance of the trace lines from the centre of each color-cell is proportional to the Gini index.

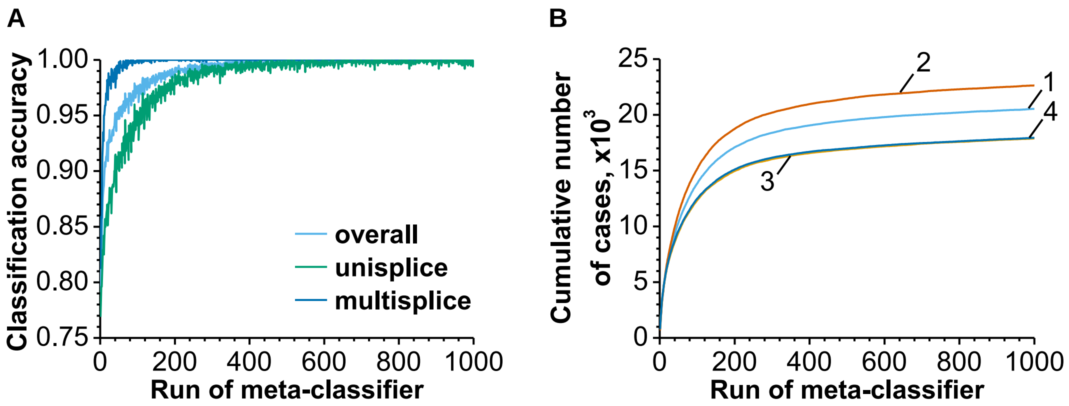


**Supplementary Figure S7.** Iterative removing of misclassified splice sites leads to a high accuracy of classification. (A) Improved classification accuracy of 5' splice sites from the siMM treated Kasumi-1 cells after multiple rounds of the random forest classifier runs and removing misclassified cases. (B) The cumulative number of misclassified cases after multiple rounds of the random forest classifier runs. Lines 1 and 2 represent 5' splice sites from the siMM and siRR treated leukemia cells, respectively. Similarly, lines 3 and 4 show 3' splice sites from the siMM and siRR treated leukemia cells, respectively.

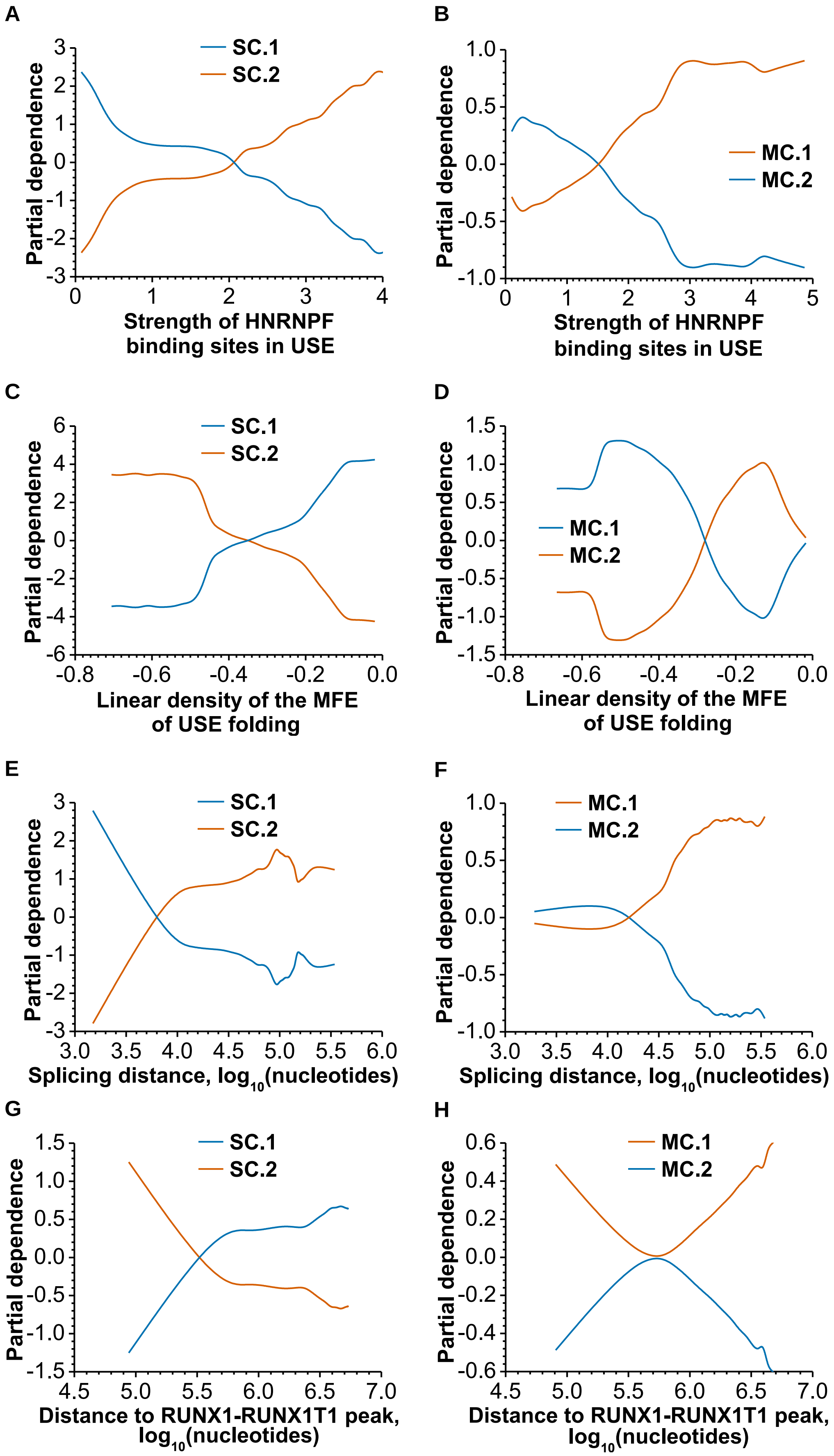


**Supplementary Figure S8.** The effects of different values of the most important features on the class probability of 5’ splice sites from the siMM treated Kasumi-1 cells dataset. Each partial dependence plot gives a graphical depiction of the adjusted effect of a given feature on the class probability in the context of the whole set of other features. Plots are based on results from 100 independent runs of the random forest based meta-classifier and 1000 classification trees per random forest per run.

(RAGE) through G-rich cis-elements and heterogenous nuclear ribonucleoprotein H. *J. Biochem.*, **147**, 651–659.

141. Simard,M.J. and Chabot,B. (2002) SRp30c is a repressor of 3' splice site utilization. *Mol. Cell. Biol.*, **22**, 4001–4010.
142. Duning,K., Buck,F., Barnekow,A. and Kremerskothen,J. (2008) SYNCRIP, a component of dendritically localized mRNPs, binds to the translation regulator BC200 RNA. *J. Neurochem.*, **105**, 351–359.
143. Aznarez,I., Barash,Y., Shai,O., He,D., Zielenski,J., Tsui,L.-C., Parkinson,J., Frey,B.J., Rommens,J.M. and Blencowe,B.J. (2008) A systematic analysis of intronic sequences downstream of 5' splice sites reveals a widespread role for U-rich motifs and TIA1/TIAL1 proteins in alternative splicing regulation. *Genome Res.*, **18**, 1247–1258.
144. Izquierdo,J.M. and Valcárcel,J. (2007) Two isoforms of the T-cell intracellular antigen 1 (TIA-1) splicing factor display distinct splicing regulation activities. Control of TIA-1 isoform ratio by TIA-1-related protein. *J. Biol. Chem.*, **282**, 19410–19417.
145. McAlinden,A., Liang,L., Mukudai,Y., Imamura,T. and Sandell,L.J. (2007) Nuclear protein TIA-1 regulates COL2A1 alternative splicing and interacts with precursor mRNA and genomic DNA. *J. Biol. Chem.*, **282**, 24444–24454.
146. Reyes,R. and Izquierdo,J.M. (2007) The RNA-binding protein PTB exerts translational control on 3'-untranslated region of the mRNA for the ATP synthase beta-subunit. *Biochem. Biophys. Res. Commun.*, **357**, 1107–1112.
147. Wang,I., Hennig,J., Jagtap,P.K.A., Sonntag,M., Valcárcel,J. and Sattler,M. (2014) Structure, dynamics and RNA binding of the multi-domain splicing factor TIA-1. *Nucleic Acids Res.*, **42**, 5949–5966.
148. Fraser,D.J., Phillips,A.O., Zhang,X., van Roeyen,C.R., Muehlenberg,P., En-Nia,A. and Mertens,P.R. (2008) Y-box protein-1 controls transforming growth factor-beta1 translation in proximal tubular cells. *Kidney Int.*, **73**, 724–732.
149. Shen,Q., Fan,L. and Newburger,P.E. (2006) Nuclease sensitive element binding protein 1 associates with the selenocysteine insertion sequence and functions in mammalian selenoprotein translation. *J. Cell. Physiol.*, **207**, 775–783.
150. Wei,W.-J., Mu,S.-R., Heiner,M., Fu,X., Cao,L.-J., Gong,X.-F., Bindereif,A. and Hui,J. (2012) YB-1 binds to CAUC motifs and stimulates exon inclusion by enhancing the recruitment of U2AF to weak polypyrimidine tracts. *Nucleic Acids Res.*, **40**, 8622–8636.
151. Murray,J.I., Voelker,R.B., Henscheid,K.L., Warf,M.B. and Berglund,J.A. (2008) Identification of motifs that function in the splicing of non-canonical introns. *Genome Biol.*, **9**, R97.
152. Li,L. (2009) GADEM: a genetic algorithm guided formation of spaced dyads coupled with an EM algorithm for motif discovery. *J. Comput. Biol.*, **16**, 317–329.
153. Droit,A., Gottardo,R., Robertson,G. and Li,L. (2014). rGADEM: de novo motif discovery. R package version 2.22.0.
154. Ameur,A. (2010). BCRANK: Predicting binding site consensus from ranked DNA sequences. R package version 1.36.0.

170. Zhou,H.-L., Baraniak,A.P. and Lou,H. (2007) Role for Fox-1/Fox-2 in mediating the neuronal pathway of calcitonin/calcitonin gene-related peptide alternative RNA processing. *Mol. Cell. Biol.*, **27**, 830–841.
171. Minovitsky,S., Gee,S.L., Schokrpur,S., Dubchak,I. and Conboy,J.G. (2005) The splicing regulatory element, UGCAUG, is phylogenetically and spatially conserved in introns that flank tissue-specific alternative exons. *Nucleic Acids Res.*, **33**, 714–724.
172. Kar,A., Havlioglu,N., Tarn,W.-Y. and Wu,J.Y. (2006) RBM4 interacts with an intronic element and stimulates tau exon 10 inclusion. *J. Biol. Chem.*, **281**, 24479–24488.
173. Lin,J.-C. and Tarn,W.-Y. (2005) Exon selection in alpha-tropomyosin mRNA is regulated by the antagonistic action of RBM4 and PTB. *Mol. Cell. Biol.*, **25**, 10111–10121.
174. Královicová,J., Houngninou-Molango,S., Krämer,A. and Vorechovsky,I. (2004) Branch site haplotypes that control alternative splicing. *Hum. Mol. Genet.*, **13**, 3189–3202.
175. Tacke,R., Tohyama,M., Ogawa,S. and Manley,J.L. (1998) Human Tra2 proteins are sequence-specific activators of pre-mRNA splicing. *Cell*, **93**, 139–148.
176. Castle,J.C., Zhang,C., Shah,J.K., Kulkarni,A.V., Kalsotra,A., Cooper,T.A. and Johnson,J.M. (2008) Expression of 24,426 human alternative splicing events and predicted cis regulation in 48 tissues and cell lines. *Nat. Genet.*, **40**, 1416–1425.
177. Culler,S.J., Hoff,K.G., Voelker,R.B., Berglund,J.A. and Smolke,C.D. (2010) Functional selection and systematic analysis of intronic splicing elements identify active sequence motifs and associated splicing factors. *Nucleic Acids Res.*, **38**, 5152–5165.
178. Das,D., Clark,T.A., Schweitzer,A., Yamamoto,M., Marr,H., Arribere,J., Minovitsky,S., Poliakov,A., Dubchak,I., Blume,J.E., *et al.* (2007) A correlation with exon expression approach to identify cis-regulatory elements for tissue-specific alternative splicing. *Nucleic Acids Res.*, **35**, 4845–4857.
179. Wang,Y., Ma,M., Xiao,X. and Wang,Z. (2012) Intronic splicing enhancers, cognate splicing factors and context-dependent regulation rules. *Nat. Struct. Mol. Biol.*, **19**, 1044–1052.
180. Wang,Y., Xiao,X., Zhang,J., Choudhury,R., Robertson,A., Li,K., Ma,M., Burge,C.B. and Wang,Z. (2013) A complex network of factors with overlapping affinities represses splicing through intronic elements. *Nat. Struct. Mol. Biol.*, **20**, 36–45.
181. Yeo,G.W., Van Nostrand,E.L., Nostrand,E.L.V. and Liang,T.Y. (2007) Discovery and analysis of evolutionarily conserved intronic splicing regulatory elements. *PLoS Genet.*, **3**, e85.
182. Banerjee,H., Rahn,A., Gawande,B., Guth,S., Valcarcel,J. and Singh,R. (2004) The conserved RNA recognition motif 3 of U2 snRNA auxiliary factor (U2AF 65) is essential in vivo but dispensable for activity in vitro. *RNA*, **10**, 240–253.
183. Mercer,T.R., Clark,M.B., Andersen,S.B., Brunck,M.E., Haerty,W., Crawford,J., Taft,R.J., Nielsen,L.K., Dinger,M.E. and Mattick,J.S. (2015) Genome-wide discovery of human splicing branchpoints. *Genome Res.*, **25**, 290–303.
184. Stojnic,R. and Diez,D. (2015). PWMEnrich: PWM enrichment analysis. R package version 4.10.0.
185. Lorenz,R., Bernhart,S.H., Höner Zu Siederdissen,C., Tafer,H., Flamm,C., Stadler,P.F. and Hofacker,I.L. (2011) ViennaRNA Package 2.0. *Algorithms Mol Biol*, **6**, 26.
